# Computational Signatures of Pain Chronification: Duration-Dependent Decision-Making Shifts Across Acute and Chronic Pain

**DOI:** 10.1101/2025.07.09.663993

**Authors:** Chad C. Williams, Lucy L.W. Owen, Chloe Z. Gunsilius, Matthew R. Nassar, Frederike H. Petzschner

**Author notes:** These authors contributed equally to this work (Chad C. Williams, Lucy L.W. Owen).

## Abstract

Chronic pain is often characterized by the entrenchment of maladaptive behaviors — avoidance, inactivity, and hypervigilance — that outlast tissue damage. A central but mechanistically underspecified question is how behaviors acquired during acute pain become locked in as pain chronifies. Influential accounts, including the fear-avoidance model, suggest that behavioral patterns learned in a specific context — such as avoidance during acute injury — overgeneralize to situations where they are no longer advantageous. Whether this reflects altered learning or changes in how learned information guides decisions has remained unclear. To address this, we administered a probabilistic reinforcement learning task to 239 individuals with chronic pain, acute pain, or no pain, designed to dissociate two decision strategies: reliance on recent reinforcement history — what was reinforced in a prior context — versus global expected value — the objective worth of an option regardless of context. Learning performance was comparable across all groups, confirming intact associative learning. However, groups differed significantly in decision-making: individuals with chronic pain favored options with stronger context-dependent reinforcement history over those with higher global expected value; however, those without pain showed no preference for one over the other, with the acute pain group showing an intermediate pattern. Computational modeling confirmed this, with chronic pain patients showing significantly reduced weighting toward global expected value. Critically, this shift tracked pain duration rather than intensity — suggesting prolonged pain exposure gradually biases decisions away from global expected value towards a context-dependent history. These findings offer a computational explanation for behavioral persistence in chronic pain, and this duration-dependent shift, already evident in the acute pain group, may represent an early cognitive signature of chronification.

## Introduction

Pain is among the most powerful teaching signals available to an organism - it signals tissue damage and drives behaviors that protect the body. But when pain persists beyond the normal healing period, typically three to six months, it becomes chronic and often maladaptive. Chronic pain affects approximately 30% of adults globally and is a leading cause of long-term disability [1–3], yet the mechanisms driving its persistence remain poorly understood.

A compelling account is offered by the fear-avoidance model, which proposes that chronic pain is maintained through maladaptive reinforcement cycles [4, 5]. Behaviors that reduce pain in the short term - rest, withdrawal, hypervigilance - are reinforced during acute injury and can persist long after tissue healing, even when they hinder recovery [6, 7]. Yet the model leaves a critical question unanswered: does chronic pain alter learning itself - the ability to acquire new associations - or how learned information guides decisions? This distinction determines where maladaptive behavior originates and where intervention should be targeted.

Computational models of reinforcement learning provide a framework to answer this question. Decision-making relies on a balance between two processes [8–11]: learning from recent reinforcement history - what was rewarded or punished in a specific context - and forming global expected value representations that summarize the long-term worth of an option regardless of context [8, 10, 12–15]. We hypothesize that chronic pain disrupts this balance, biasing decisions toward recent reinforcement history at the expense of global expected value - a shift that would explain why avoidance behaviors acquired during acute pain persist even when no longer adaptive. Supporting this view, corticostriatal circuits linking the nucleus accumbens and ventromedial prefrontal cortex - regions central to this balance in value-based decision-making [8, 10] - predict the transition from acute to chronic pain [16, 17], and individuals at risk of chronification show altered striatal responses to reward prediction errors [18].

To test this hypothesis, we administered a probabilistic reinforcement learning task to 239 individuals with chronic pain, acute pain, or no pain [19–21]. The task quantifies the degree to which individual decisions are driven by recent reinforcement history versus global expected value by comparing choices between options learned in separate contexts (punishment-avoidance and reward-seeking). We applied computational modeling to estimate each participant’s relative weighting of these two strategies at the level of individual choices.

We expected chronic pain to be associated with reduced weighting of global expected value in favor of recent reinforcement history, which would provide a computationally specified mechanism for the behavioral persistence described by the fear-avoidance model and raise the possibility that individual decision-making profiles during acute and sub-acute stages could serve as early markers of chronification risk.

## Results

To investigate how reward-based decision making varies among individuals in different pain states, we adapted a well-established probabilistic two-armed bandit task previously used to dissociate whether participants’ choices reflect absolute expected value integration or are instead shaped by local context associations and reinforcement dynamics [20, 21]. In our study, a modified version of this task was administered to 239 individuals experiencing either chronic (*n* = 78), acute (*n* = 53), or no pain (*n* = 108), allowing us to systematically examine how pain states influence learning and value-based choice behavior (Figure 1a).

**Fig. 1.**
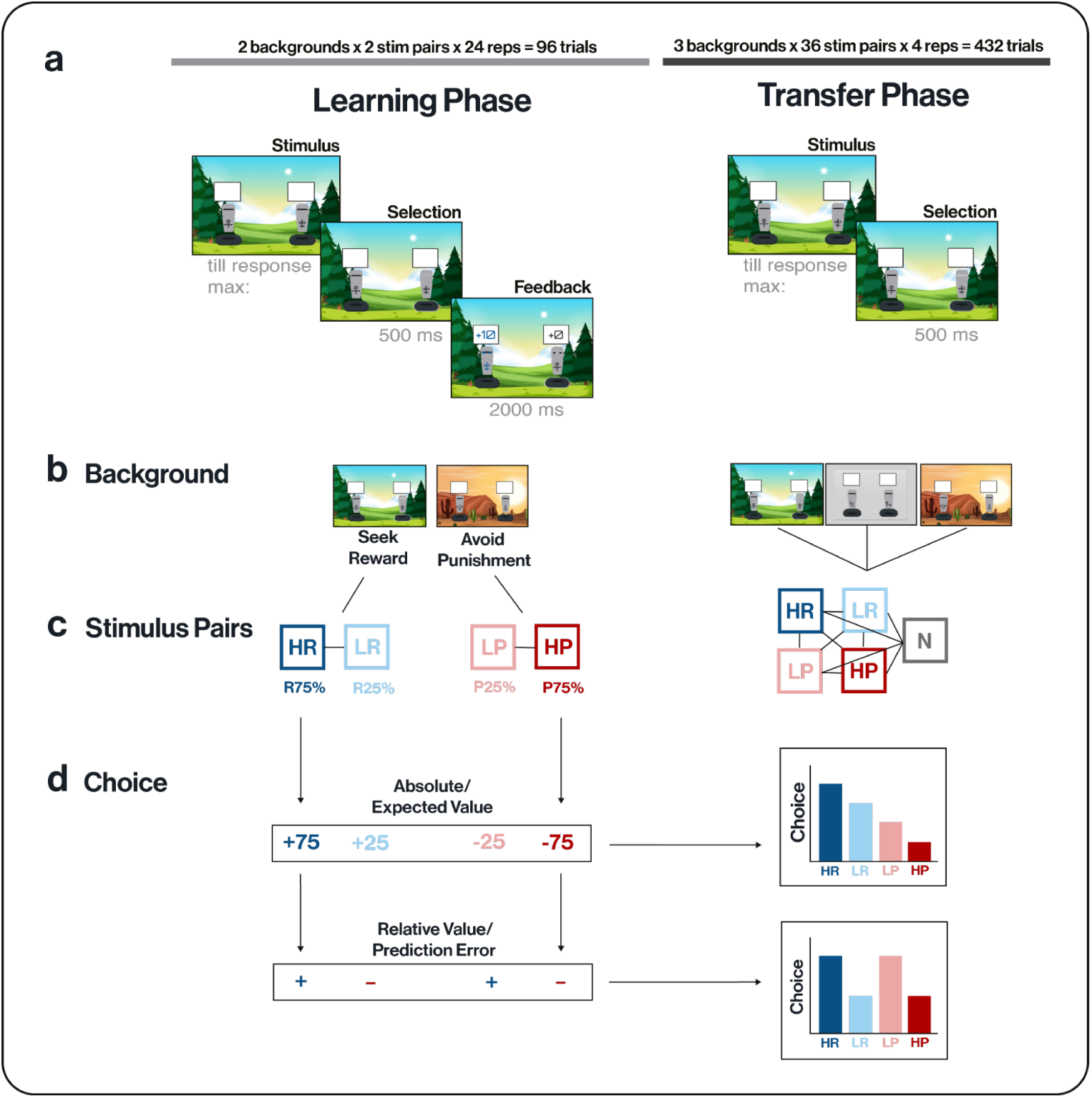
Task structure. a. Trial Scheme: Each trial began with the presentation of a fixed stimulus pair (artificial knights dissociable by the symbol on their harness) and a contextual background image (forest or desert). Participants selected one of the two stimuli (knights) and received feedback indicating either reward or punishment, depending on the context. During the learning phase, feedback was provided after every choice for both the chosen and unchosen options, whereas in the transfer phase, participants made choices between novel pairings of previously learned stimuli without receiving feedback. b. Context Backgrounds: In the learning phase, each background image was uniquely associated with a stimulus pair type: i.e., the desert background signaled a reward-seeking context (2 unique pairs of low-reward rate [LR] and high-reward rate [HR] stimuli), while the forest background signaled a punishment-avoidance context (2 unique pairs of low-punishment rate [LP] and high-punishment rate [HP] stimuli). In the transfer phase, all possible pairwise combinations of the four stimulus types (HR, LR, LP, HP) and a never-seen novel stimulus (N) were presented across three background conditions: the original reward background, the original punishment background, and a neutral grey background. This design allowed us to assess whether contextual cues provided by visual background or by the original stimulus pairings continued to influence decision-making. c. Stimulus Pairs: During the learning phase, stimuli were paired exclusively within valence domains: HR and LR were always presented together in the reward context, and LP and HP were always paired in the punishment context (2 stimulus pairs per category). HR stimuli yielded rewards on 75% of trials, LR on 25%, and otherwise a neutral outcome. Conversely, HP stimuli resulted in losses on 75% of trials, while LP stimuli resulted in losses on 25% of trials. During the transfer phase, all previously experienced stimuli (2x HR, LR, LP, HP) and a previously unseen novel stimulus (N) were paired in combinations with one another, resulting in 36 possible unique stimulus pairings. d. Potential Choice Strategies: If participants based their choices on absolute expected values, preferences should follow the order HR > LR > LP > HP. However, if decision-making relied on context-specific relative value or the frequency of positive prediction errors (PEs) during learning, LP stimuli—although objectively worse than LR—might be preferred due to their relatively advantageous status within their original context and frequent positive prediction errors. The plot illustrates predicted choice probabilities for these competing strategies across critical stimulus comparisons. Abbreviations: HR – high reward rate (75% reward), LR – low reward rate (25% reward), LP – low punishment rate (25% loss), HP – high punishment rate (75% loss).

During the initial learning phase of the task, participants repeatedly selected between fixed pairs of stimuli presented in either a reward context or a punishment- avoidance context. Context was signalled in two ways: a background image indicating either a forest or a desert scene, and the unique pairing of two stimuli (Figure 1b,c). Within each context, one stimulus was objectively more advantageous than the other. Specifically, participants learned to choose between a stimulus with a high probability of reward (75%, high reward rate, HR) and a low probability of reward (25%, low reward rate, LR) in the reward seeking context, or between a low-probability punishment stimulus (25% loss, low punishment rate, LP) and a high-probability punishment stimulus (75% loss, high punishment rate, HP) in the punishment avoidance context. These four stimulus types differed both in their absolute expected value and their relative standing within their local context (Figure 1d). During learning, participants received outcome feedback after each choice for both the chosen and unchosen stimuli. However, only the chosen option resulted in actual gains or losses.

After learning, participants entered a transfer phase in which all stimuli were recombined into novel pairings and presented without feedback. Critically, this phase was designed to reveal which aspects of the learning phase shaped choice behavior. Participants were now required to make decisions between stimuli that had never been directly compared, for instance, a high-reward rate stimulus versus a low-punishment rate stimulus.

These novel pairings were constructed to dissociate two distinct decision-making strategies. One strategy involves selecting the option with the highest absolute expected value—a global representation learned across all contexts based on outcome probabilities. The alternative strategy relies on context-specific learning signals, such as the relative value of a stimulus within its original pair, i.e., based on the frequency of positive prediction errors it elicited during learning. For example, while the low-punishment rate (LP) stimulus had a negative absolute expected value, it was the relatively better option in its original punishment context and consistently produced positive prediction errors. While the low-reward rate stimulus (LR) had a positive absolute expected value, it was the relatively worse option in its original context and consistently produced negative prediction errors. As a result, whereas participants who rely on absolute values would prefer the LR stimulus over the LP stimulus, those whose decisions are guided more by relative value or prediction-error dynamics may instead favor the LP stimulus (Figure 1d).

### Learning Performance Does Not Differ Across Pain Groups, but Chronic Pain is Associated with Slower Response Times

Participants across all three groups—chronic pain, acute pain, and no pain—demonstrated comparable learning performance during the initial phase of the task (Figure 2a). Accuracy improved with repeated exposure to stimulus pairs, and critically, no group differences emerged in the ability to learn optimal actions within either reward-seeking or punishment-avoidance contexts. A generalized linear mixed-effects model (GLMM) revealed a significant main effect of time (*p* < .0001), reflecting overall learning progression. However, there were no significant main effects of group (*p* = .9192) or context (reward seeking vs. punishment avoidance; *p* = .1406), nor evidence for interactions involving group and context (*ps* > .14), except for a modest group × time interaction (*p* = .0037), suggesting slight variation in learning curves over time (Supplementary Table S1). Planned comparisons confirmed the absence of group differences: neither the chronic pain group versus no pain (*p* = .4730, *d* = –0.11) nor chronic versus acute pain (*p* = .7991, *d* = –0.05) differed significantly. No effects were found when comparing reward and punishment contexts across groups (*ps* > .05; Supplementary Table S1).

**Fig. 2.**
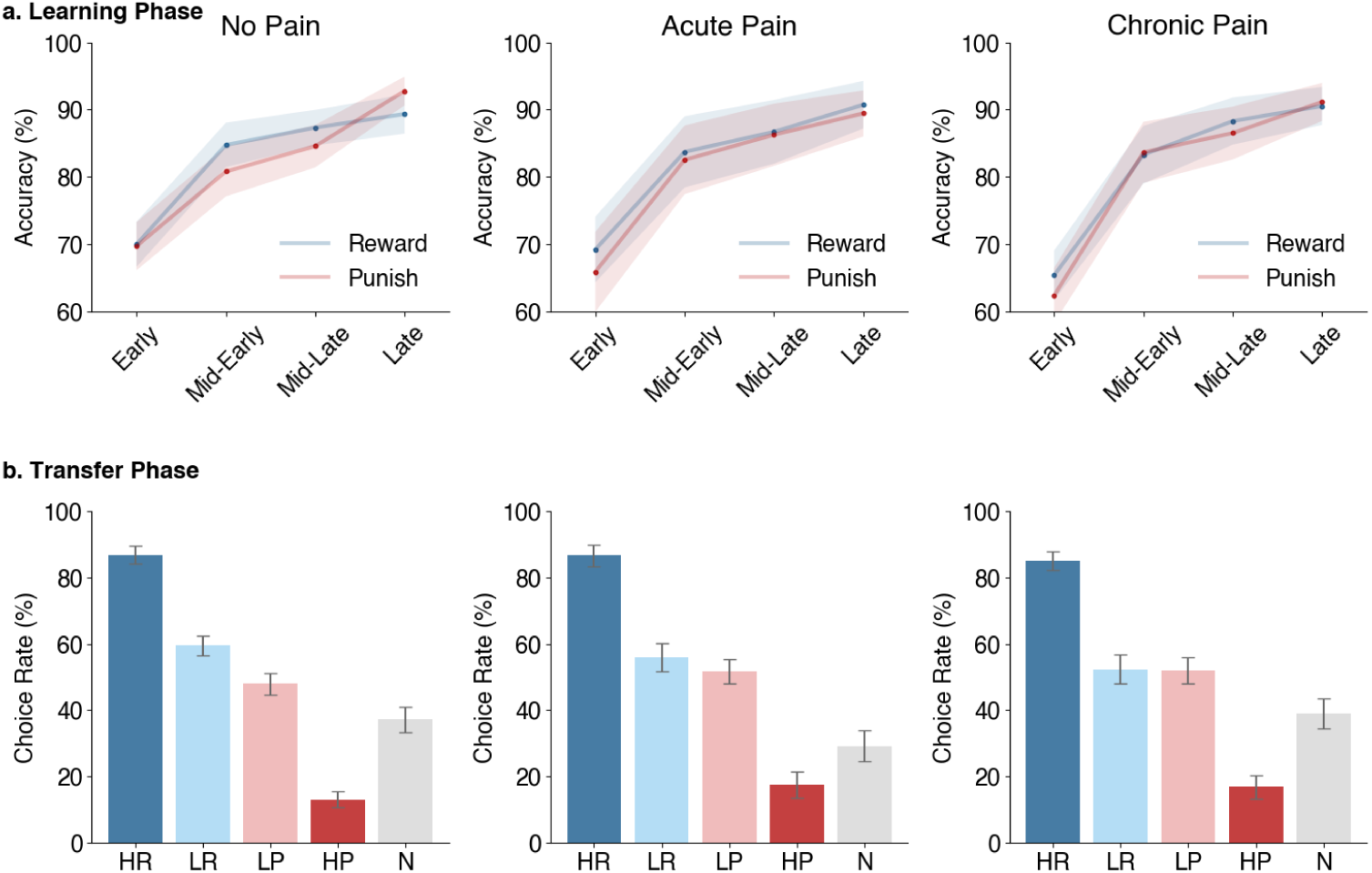
Empirical findings of learning accuracy and transfer choice rates. a. Learning Phase: Behavioral performance across binned learning trials for the reward and punishment contexts for each group. Shaded regions represent 95% confidence intervals. b. Transfer Phase: Choice rates for each stimulus type during transfer trials for each group. Choice rate is computed as the percentage of times a stimulus type was chosen, given the number of times it was presented. Bar plots show the mean and 95% confidence intervals of the choice rate for each stimulus type across participants within each group. Abbreviations: HR – high reward rate (75% reward), LR – low reward rate (25% reward), LP – low punishment rate (25% loss), HP – high punishment rate (75% loss), N – novel stimulus.

By contrast, response times differed significantly across groups. Participants with chronic pain responded more slowly than both pain-free individuals and those with acute pain, despite equivalent accuracy (Supplementary Figure S1a). The GLMM revealed significant main effects of group (*p* = .0441), context (*p* < .0001), and time (*p* < .0001), as well as significant group × context (*p* = .0056) and group × time (*p* = .0007) interactions (Supplementary Table S2). Planned comparisons showed slower responses in the chronic pain group relative to both the no pain (chronic > no pain: *p* = .0020, *d* = 0.47) and acute pain (chronic > acute: *p* = .0224, *d* = 0.41) groups. These differences held across both reward (*p* = .0011, *d* = 0.51) and punishment (*p* = .0083, *d* = 0.40) contexts, although no differences were found in post hoc comparisons of reaction times between the acute and no pain groups (*p* = .9047, *d* = –0.07). In addition, overall reaction times were slower in the punishment than in the reward context and decreased across time.

Together, these findings support that basic learning performance is intact in individuals with both acute and chronic pain, and that the ability to learn to seek rewards or avoid punishments does not differ between pain groups. However, individuals with chronic pain exhibited significantly slower response times, suggesting alterations in decision speed rather than learning capacity.

### Pain Groups Differ in Reward-Based Decision Making During the Transfer Phase

There are at least two distinct ways participants could have learned to make accurate choices during the learning phase: by encoding the relative values of options within each context (i.e., learning which option was better in a given pair), or by computing the overall expected value of individual stimuli across trials. The transfer phase was designed to disentangle these strategies. In this phase, previously learned stimuli—along with a novel stimulus—were presented in new pairings across various background contexts, without feedback (Figure 1). Because options were now combined across contexts that differed in their overall reward structure, participants’ choices revealed whether they had formed absolute value estimates (i.e. HR > LR > LP > HP) or relied on context-specific reinforcement (i.e. HR and LP > LR and HP) (Figure 1d). Thus the transfer phase allowed us to test our central hypothesis that valuation strategies differ across groups in different pain states.

Indeed, we find that participants with chronic pain relied more heavily on context-specific reinforcement history than on globally integrated expected value representations during the transfer phase. Choice behavior during the transfer phase varied across stimulus types, which significantly interacted with group identity. A generalized linear mixed-effects model (GLMM) revealed a robust main effect of stimulus type (HR, LR, LP, HP, Novel [N]; *p* < .0001), no main effect of group (*p* = .1408), but a significant group × stimulus type interaction (*p* = .0002) (Figures 2b and 3; Supplementary Figure S2; Table 1). Planned comparisons indicated that the chronic pain group differed significantly in their choice preferences from both the no pain group (*p* = .0136, *d* = –0.37) and the acute pain group (*p* = .0420, *d* = –0.37) (Table 1).

**Fig. 3.**
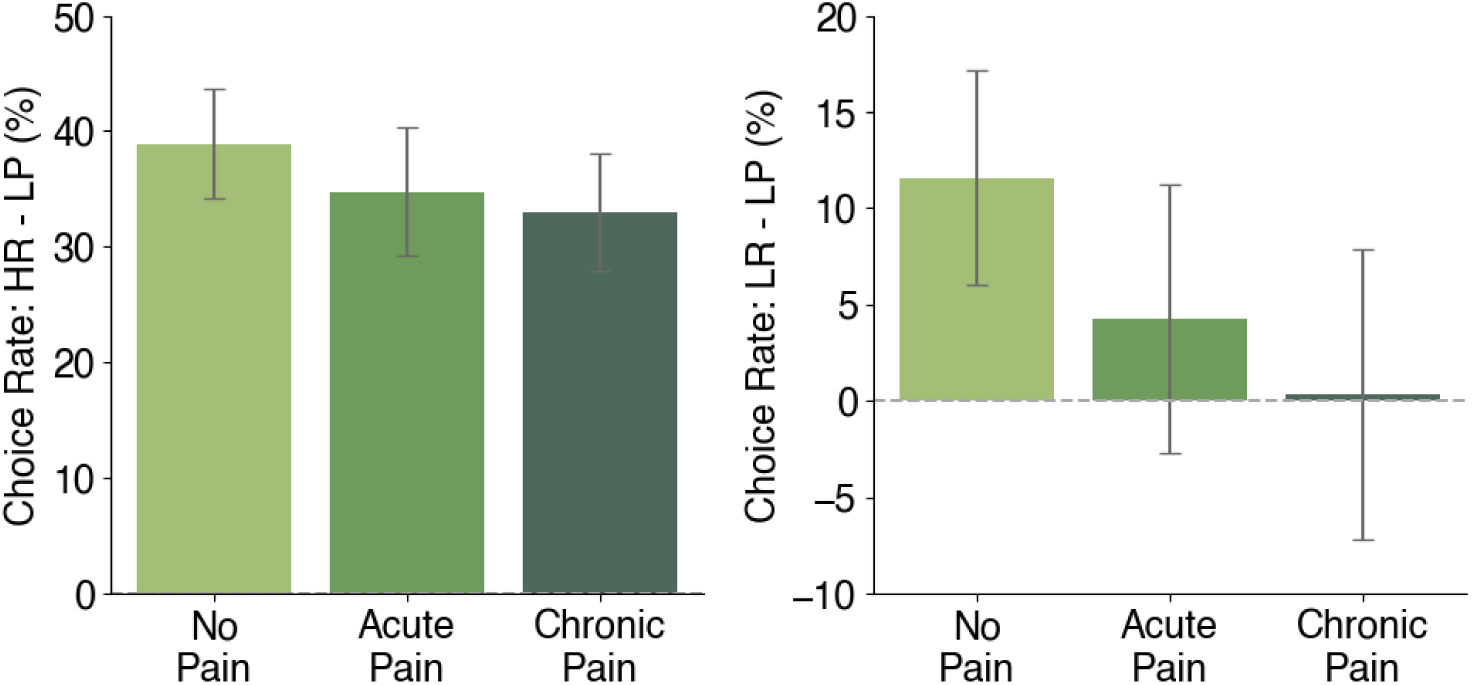
Choice rate differences for critical analyses assessing valuation strategies. Difference in choice rates (%) comparing the high reward (HR) with the low punish (LP) stimulus type and the low reward (LR) with the low punish (LP) stimulus type across the groups. Choice rate is here computed as the number of times that the stimulus type was selected, given the number of times it was presented in its pair. This was computed for each stimulus in the pair (e.g., HR and LP for the left plot and LR and LP for the right plot), and the difference of these is plotted here. Bar plots show the mean and 95% confidence intervals of the difference in choice rates.

**Table 1.**
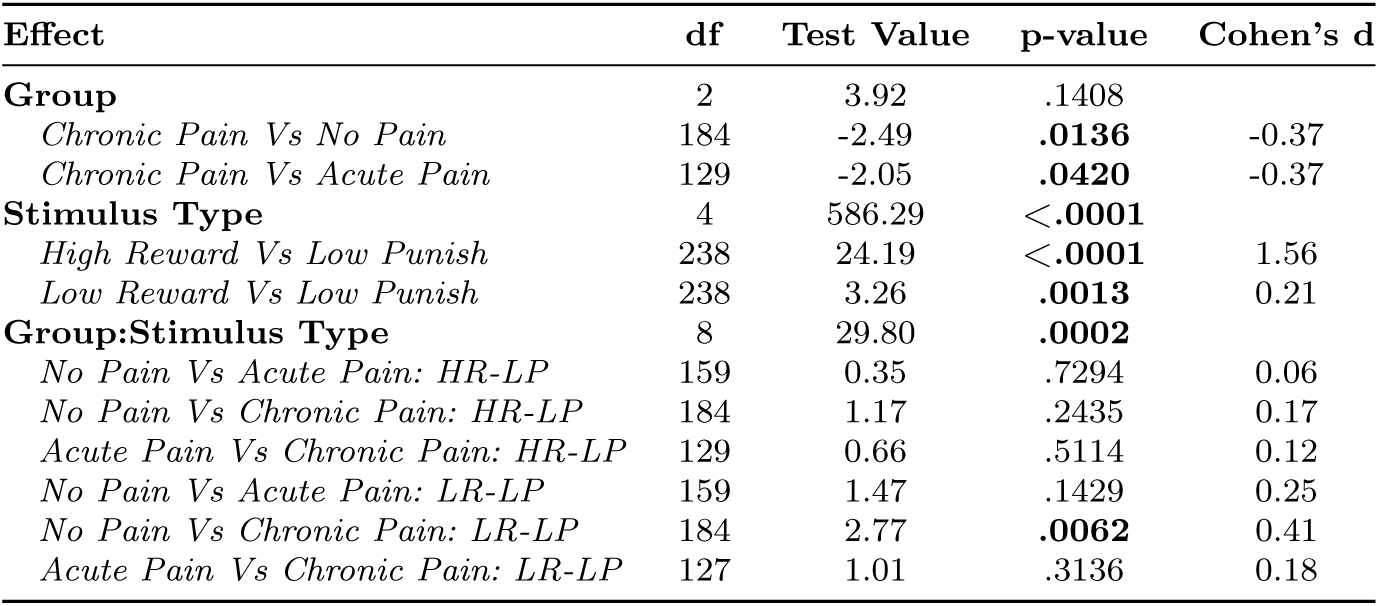
Analysis of Choice Rates During the Transfer Phase. Statistical outcomes of GLMMs (**bold effects**) and planned comparisons (indented and *italicized eflects*) for choice rates in the transfer phase. Test value represents *χ*^2^ for GLMM effects and t-values for planned comparisons. The planned comparisons of the group x stimulus type interaction use slightly different data than the GLMMs and the other planned comparisons. Specifically, these assess choice rates for each pair (e.g., HR vs LP) only when presented in that pair (rather than choice rates across all pairs). **Bold p-values** indicate statistically significant effects at *α* = .05.

These effects remained robust when controlling for background context (i.e., the visual background image, Supplementary Table S3). This lack of influence from the visual context images suggests that the observed group differences in valuation strategies cannot be explained by differential incorporation of contextual features into the state representation and suggests that participants’ decisions were driven by previously encountered stimulus evaluations rather than contextual cues. Moreover, performance differences were consistent across the first and second halves of the transfer phase, suggesting that valuation strategies remained stable over time and were not confounded by fatigue or task disengagement.

Reaction time patterns during the transfer phase paralleled the main effects in choice behavior. While there was no main group effect on overall reaction times (*p* = .8893), there was a significant main effect of stimulus type (*p* < .0001) and a group × stimulus type interaction (*p* < .0001; see Supplementary Figure S1b and Supplementary Table S4).

Overall, these results indicate that the chronic pain group adopted a different valuation strategy during the transfer phase—favoring context-specific reinforcement over integrated value representations—despite similar learning performance across groups.

### Choices in Chronic Pain Are Biased Towards Context-Dependent Reinforcement History

To determine whether distinct valuation strategies drove the observed group differences during transfer, we focused on stimulus pairings that dissociate absolute expected value from context-specific reinforcement history. Specifically, we analyzed choices and reaction times for HR vs. LP and LR vs. LP stimulus pairs.

The HR vs. LP contrast compares two stimuli that were both the better option within their original learning contexts and thus share similar reinforcement histories (i.e., frequent positive prediction errors). However, they differ substantially in absolute value: HR carries a positive expected value, whereas LP has a negative one (Figure 1d). A strong preference for HR over LP reflects sensitivity to global expected value independent of reinforcement history.

Across groups, participants showed a clear preference for HR over LP, indicating a shared sensitivity to absolute expected value when reinforcement history was similar. Across groups, we found no significant differences in HR vs. LP choice rates, suggesting that all participants relied to some extent on absolute expected value when reinforcement history was similar. Specifically, planned comparisons of choice behavior did not differ between the no pain and acute pain groups (*p* = .7294, *d* = 0.06), the acute and chronic pain groups (*p* = .5114, *d* = 0.12), or the no pain and chronic pain groups (*p* = .2435, *d* = 0.17) (Table 1). Similarly, reaction time data of the HR vs. LP contrast revealed no group differences in choice speed between the no pain and acute pain groups (*p* = .3361, *d* = −0.18), the no pain and chronic pain groups (*p* = .2966, *d* = −0.17), and the acute and chronic pain groups (*p* = .9095, *d* = 0.02) (Supplementary Table S4).

The LR vs. LP comparison contrasts opposing reinforcement histories and value profiles. LR has a higher expected value but was the worst option within its original reward context, typically associated with negative prediction errors. In contrast, LP has a lower expected value but was contextually positively reinforced. Preference for LR over LP reflects sensitivity to global expected value; preference for LP over LR reflects reliance on context-specific reinforcement.

Here, group differences in choices were evident. Participants in the chronic pain group were significantly less likely to prefer LR over LP compared to the no pain group (chronic < no pain: *p* = .0062, *d* = −0.41), indicating greater reliance on reinforcement history. The acute pain group did not differ significantly from either comparison group (acute vs. no pain: *p* = .1429, *d* = −0.25, acute vs. chronic pain: *p* = .3136, *d* = 0.18), but had choice rates that were numerically between the chronic pain and no pain group (Figure 3). Post hoc analyses showed that only the no pain group reliably preferred LR over LP (*p* < .0001, *d* = 0.44), while both pain groups treated the options similarly (acute: *p* = .3693, *d* = 0.22; chronic: *p* = 1.0000, *d* = 0.02). In contrast, there were no effects of reaction times when comparing the LR to LP stimulus types (Supplementary Table S4). Specifically, there was no difference between the no pain and the acute pain groups (*p* = .3052, *d* = −0.18), the no pain and chronic pain groups (*p* = .1400, *d* = −0.23), or the acute pain and chronic pain groups (*p* = .7528, *d* = −0.06).

Together, these findings indicate a progressive shift from expected value–driven decision-making in pain-free individuals toward a greater influence of reinforcement history–based valuation in individuals with chronic pain, with the acute pain group occupying a transitional position between the two. Furthermore, this shift occurred independently of response time differences, thus reflecting a genuine shift in valuation strategies rather than changes in processing speed.

### Decision-Making Across Pain States Is Best Explained by a Mixture Model of Expected Value and Reinforcement History

While behavioral analyses identified group-level differences in choice patterns, computational modeling provides a more granular account of the cognitive mechanisms that shape these decisions. To this end, we fit five reinforcement learning models to participants’ trial-by-trial choices across both the learning and transfer phases. These models systematically varied in how they weighted global expected value versus local context-specific reinforcement history (see Methods for full specifications and Supplementary Information for model validations).

The Q-Learning model captures decision-making based on the integration of past outcomes into an absolute expected value for each option (pure expected value strategy) [22, 23]. By contrast, the Actor-Critic model distinguishes between state and action learning, emphasizing recent reinforcement signals without forming stable value representations (pure reinforcement history strategy) [23, 24]. The remaining three models implement mixture strategies that combine global and local learning mechanisms typically captured by a single model parameter (mixture models). The Hybrid model uses a weighted combination of Q-Learning and Actor-Critic mechanisms [19, 20]. The Relative model integrates Q-values with within-context value normalization, capturing the dynamic scaling of preferences based on local context [21]. Finally, the Advantage model, which we introduce here, simplifies this approach by incorporating both absolute Q-values and a context-dependent advantage term that reflects reward comparisons within each pair. Thus, our model set contained one model that learned global values (Q-Learning), one that learned relative values (Actor-Critic), as well as three that enabled a mixture of global and relative value learning (Hybrid, Relative, Advantage).

Model comparison using the Bayesian Information Criterion (BIC) revealed that the Advantage model performed best across all three groups (Figure 4, see Supplementary Table S5 for fitted values). This model dynamically integrates global expected value estimates with recent context-specific reinforcement, allowing it to flexibly capture shifts in valuation strategy. Posterior predictive checks indicate that the model replicates the key behavior findings for both the learning and transfer phases of the task across all groups (Figure 5a,b; Supplementary Tables S6 and S7). The following results focus on group differences in key parameters from the Advantage model (but see Supplementary Information for results from the Relative and Hybrid models).

**Fig. 4.**
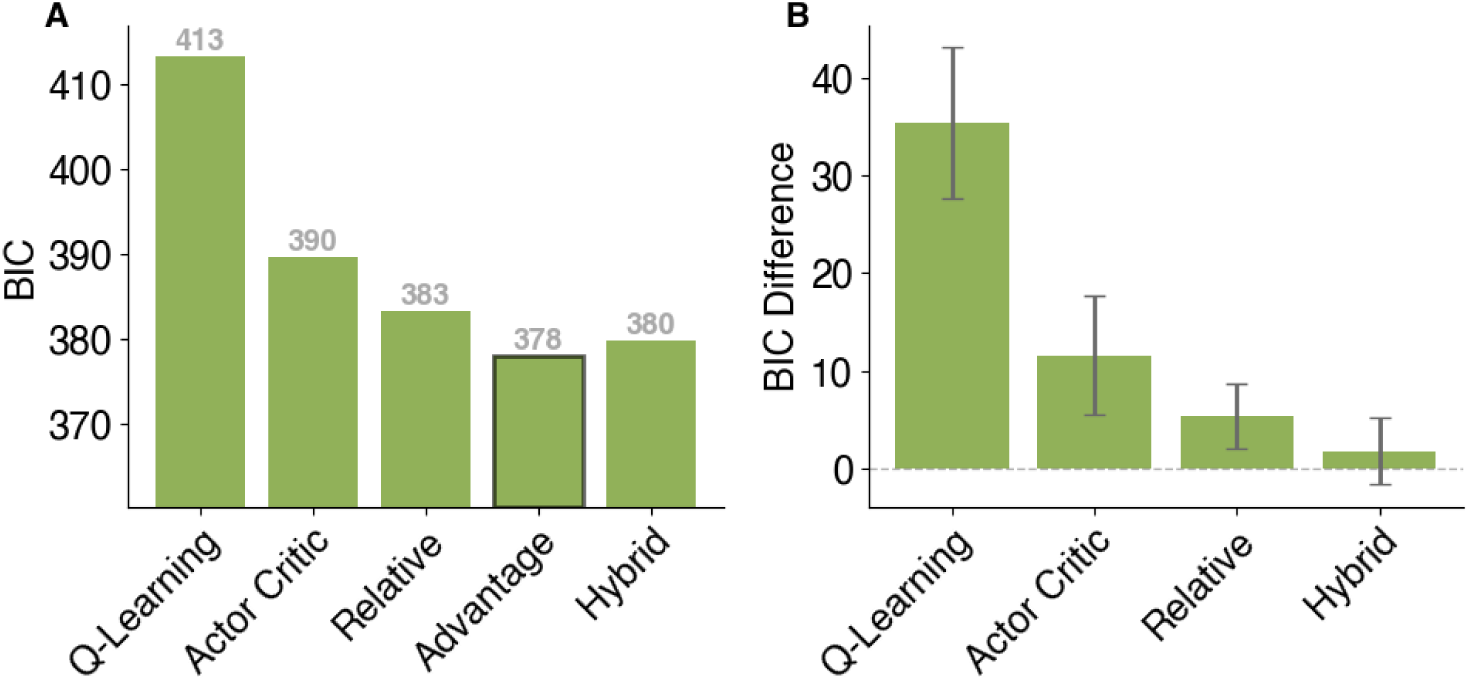
Model selection using Bayesian information criterion (BIC). A: Grand averaged BIC metrics for each model across all participants. Lower values indicate better model fit. The black border indicates the best model. B: Difference in grand average BIC values for each model relative to the best model. Error bars represent 95% confidence intervals.

**Fig. 5.**
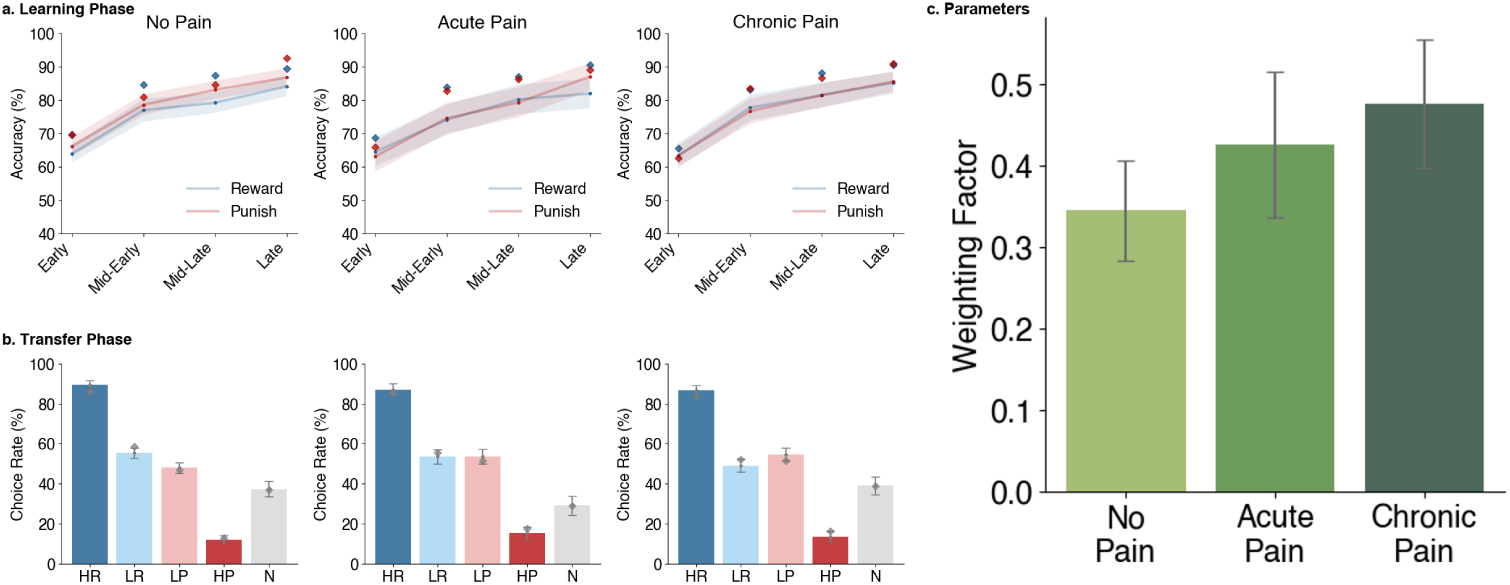
Posterior Predictive checks of the best-fitted computational model and fitted parameters. Model simulations of the best-fitted model, the Advantage model. During the fitting procedure, each participant was fitted ten times, and the best of these runs was selected to determine the participant’s fitted parameters. We then used each model to simulate a single dataset per participant using these fitted parameters. Thus, these data represent the same sample size as our empirical data. a. Learning Phase: Model performance across binned learning trials for the reward and punishment contexts for each group. Shaded regions represent 95% confidence intervals. Blue and red diamonds indicate empirical means of participant accuracy. b. Transfer Phase: Choice rates for each stimulus type during transfer trials for each group. Choice rate is computed as the percentage of times a stimulus type was chosen, given the number of times it was presented. Bar plots show the mean and 95% confidence intervals of the choice rate for each stimulus type across participants within each group. Grey diamonds indicate empirical means of participant choice rates. Abbreviations: HR – high reward rate (75% reward), LR – low reward rate (25% reward), LP – low punishment rate (25% loss), HP – high punishment rate (75% loss), N – novel stimulus. c. Parameters: Fitted values of the significant parameter from the Advantage model. The parameter is displayed as a bar plot, showing the mean and 95% confidence intervals across participants for each group.

### Chronic Pain is Associated with Increased Weighting of Context-Dependent Relative Values in Decision-Making

To identify the mechanisms underlying group differences in choice behavior, we examined fitted parameters from the best-fitting Advantage model. Of all model parameters, only the weighting factor—which determines the relative influence of global expected value versus local context-specific reinforcement—differed significantly across groups (*p* = .0270) (Figure 5c, Supplemental Figure S3, and Table 2). Planned comparisons revealed that participants with chronic pain assigned significantly less weight to global expected value than those without pain (*p* = .0088, *d* = –0.39), indicating a stronger reliance on relative context-dependent stimulus evaluations. The acute pain group did not differ significantly from either the no pain (*p* = .1394, *d* = –0.25) or chronic pain group (*p* = .6678, *d* = 0.15), again showing an intermediate pattern.

**Table 2.** Analysis of Group Differences in Model Parameters. Main effects of between-subject ANOVAs (**bold**) and planned comparisons (indented and *italicized*) for each parameter assessing group differences. Test value represents F-values for ANOVA effects and t-values for planned comparisons. **Bold p-values** indicate statistically significant effects at *α* = .05.

| Effect | df | Test Value | p-value | Cohen's d |
| --- | --- | --- | --- | --- |
| **Factual LR** | 2, 236 | 0.13 | .8792 |  |
| *No Pain Vs Acute Pain* | 159 | 0.50 | .6143 | 0.08 |
| *No Pain Vs Chronic Pain* | 150 | 0.32 | .7516 | 0.05 |
| **Counterfactual LR** | 2, 236 | 0.04 | .9574 |  |
| *No Pain Vs Acute Pain* | 159 | 0.01 | .9897 | 0.00 |
| *No Pain Vs Chronic Pain* | 184 | 0.28 | .7809 | 0.04 |
| **Temperature** | 2, 236 | 0.11 | .8991 |  |
| *No Pain Vs Acute Pain* | 159 | -0.47 | .6424 | -0.08 |
| *No Pain Vs Chronic Pain* | 184 | -0.05 | .9590 | -0.01 |
| **Weighting Factor** | 2, 236 | 3.67 | **.0270** |  |
| *No Pain Vs Acute Pain* | 159 | -1.49 | .1394 | -0.25 |
| *No Pain Vs Chronic Pain* | 184 | -2.65 | **.0088** | -0.39 |
| **Novel Value** | 2, 236 | 1.57 | .2096 |  |
| *No Pain Vs Acute Pain* | 159 | 1.57 | .1192 | 0.26 |
| *No Pain Vs Chronic Pain* | 184 | -0.27 | .7848 | -0.04 |

This computational result mirrors the empirical differences observed in LR vs. LP comparisons, suggesting a gradual shift from expected value–based to reinforcement history–driven valuation across the pain spectrum. The weighting factor from the Advantage model was highly correlated with analogous parameters in the other mixture models (Relative and Hybrid model), suggesting that this parameter reliably captures a generalizable decision strategy across model architectures (see Supplementary Information).

No other model parameters—including learning rates for chosen and unchosen options, softmax temperature, or valuation of the novel stimulus—differed significantly between groups (*ps* > .05, Table 2), underscoring the specificity of the weighting factor as the key computational mechanism underlying group differences in decision-making.

### Pain Duration, but not Severity, is Associated with Shifts in Valuation Strategy

The observed shift from expected value–based to reinforcement history–driven decision-making across pain groups appears to be primarily driven by the duration of pain rather than its subjective severity. Across participants, the weighting parameter from the best-fitting Advantage model—reflecting the relative reliance on global expected value versus local reinforcement history—was significantly associated with self-reported pain duration (ranging from ‘I don’t have pain’ to ‘pain lasting longer than 10 years’; *ρ* = .17, *p* = .0086) but not with pain severity metrics (*ps* > .05; for example, the weighting parameter was not significantly related to the composite pain score in the no pain [*r* = −.12 *p* = .2115], chronic pain [*r* = −.18 *p* = .1893], or chronic pain groups [*r* = −.03 *p* = .7972], see Supplementary Table S8). Thus, prolonged exposure to pain, rather than its momentary severity, may gradually bias decision-making toward context-dependent reinforcement signals.

None of the other model parameters were related to pain duration (factual learning rate: *ρ* = −.02, *p* = .8175; counterfactual learning rate: *ρ* = .02, *p* = .7943; temperature: *ρ* = .05, *p* = .4833; novel value: *ρ* = .01, *p* = .9133). However, within the chronic pain group, higher levels of pain unpleasantness, interference, and composite symptom scores were positively correlated with both factual and counterfactual learning rates, as well as the softmax temperature parameter (*rs* > .23, all *ps* < .05; Supplementary Figure S4 and Supplementary Table S8). These associations were absent in the acute pain group (Supplementary Figure S5 and Supplementary Table S8), despite having the same average pain intensities as the chronic pain group (Supplementary Table S9).

Taken together, these findings suggest that while valuation strategy is modulated by the chronicity of pain, symptom severity in chronic pain may independently influence how strongly individuals update beliefs and how consistently they translate those beliefs into actions, contributing to more reactive and less stable decision-making profiles.

## Discussion

This study provides evidence that chronic pain is associated with a systematic shift in how decisions are made, rather than how associations are learned. Compared to pain-free individuals, participants with chronic pain relied more on recent reinforcement history - what was locally advantageous in a prior context - and less on globally integrated expected value. This shift was confirmed computationally: the best-fitting model, which integrates both valuation signals, revealed that individuals with chronic pain assigned significantly less weight to global expected value. Crucially, this pattern was not explained by pain intensity but by pain duration, with the acute pain group showing an intermediate profile consistent with a gradual, chronification-linked process.

These findings offer a computationally specified mechanism for one of the central puzzles in chronic pain: why maladaptive behaviors persist long after tissue healing. The fear-avoidance model [4, 5] proposes that avoidance, inactivity, and hypervigilance are initially reinforced during acute injury and become entrenched over time. Our results provide a cognitive mechanism for this entrenchment: if decisions are increasingly dominated by context-dependent reinforcement history at the expense of global expected value, then behaviors that once provided short-term relief or avoided increased pain early on will continue to be selected even after the context has changed and they are no longer adaptive. The longer someone has been in pain, the more their choices appear to be guided by what worked previously rather than what is objectively best now - a computational account of how pain-related avoidance resists extinction. The finding that this shift tracks pain duration rather than intensity has important implications. Pain intensity is a momentary state; duration reflects cumulative exposure. That the valuation shift scales with duration both across acute and chronic pain groups and within the chronic pain group suggests a gradual, experience-dependent reconfiguration of decision-making architecture rather than an acute effect of nociception. This is consistent with evidence that corticostriatal circuits - linking the nucleus accumbens and ventromedial prefrontal cortex, regions central to balancing reinforcement history and global value signals - undergo plastic changes during pain chronification [16–18, 25, 26]. The weighting parameter from our best-fitting model scaled continuously with individual pain duration, suggesting a dimensional rather than categorical shift. Notably, within the chronic pain group, pain severity independently influenced other aspects of decision-making: higher pain unpleasantness and interference were associated with faster belief updating and less consistent translation of learned values into choices - suggesting that while duration shapes the valuation strategy, severity modulates how reactively and stably that strategy is expressed.

The intermediate profile of the acute pain group is particularly notable. Although this group did not differ significantly from either the chronic or pain-free group, their pattern of performance - falling consistently between the two across both behavioral and modeling analyses - raises the possibility that the valuation shift begins during acute pain and deepens with chronification. If replicated longitudinally, individual decision-making profiles at the acute stage could serve as early cognitive markers of chronification risk after injury, informing intervention before maladaptive behavioral patterns become entrenched.

The observed valuation shift maps onto neurocomputational frameworks of decision-making that distinguish between two partially dissociable systems: a dopaminergic, striatum-based system that updates choices based on prediction errors (captured by Actor-Critic learning, which reinforces options that were recently advantageous within a given context), and a prefrontal system centered in the orbitofrontal cortex and vmPFC that encodes global expected value across contexts (as in Q-Learning; [8, 10]). Individuals with chronic pain appear to rely more heavily on the former at the expense of the latter - consistent with Löffler et al. [18], who found heightened prediction error signals in the nucleus accumbens and disrupted corticostriatal dynamics in individuals who subsequently developed chronic pain. A mixture model best captured behavior across all three groups [19–21], with its weighting parameter - reflecting the relative influence of global versus context-specific reinforcement signals - scaling systematically with pain duration. This approach allows us to quantify the balance between these valuation systems at the level of individual behavior, contributing to a growing literature applying computational modeling to disentangle latent decision-making processes in clinical populations [27–38].

### Limitations

Several limitations warrant consideration. First, pain group membership was based on self-reported duration without clinical verification. Participants were recruited online via Prolific and no medical records, diagnostic imaging, or clinician assessments were obtained. While our ICD-11-informed self-report criteria provide a principled basis for group classification in a large cohort, the findings require replication in clinically well-characterized samples.

Second, we did not measure clinical avoidance behavior in real-world settings. The connection between the observed valuation shift and clinical avoidance - as indexed by measures such as the Tampa Scale of Kinesiophobia or the Pain Anxiety Symptoms Scale - remains to be tested, with the prediction that the computational weighting parameter should correlate with avoidance behavior, fear of movement, or functional disability.

Third, psychological comorbidities commonly associated with chronic pain - including depression, anxiety, and pain catastrophizing - were not systematically controlled for in the current sample. These factors are known to influence reward-based decision-making [27, 39] and may contribute to the observed group differences in valuation strategy. Future work should disentangle the specific contributions of pain chronicity and affective comorbidities to decision-making profiles.

Fourth, our sample was not stratified by chronic pain diagnosis or etiology. Chronic pain is a heterogeneous condition spanning inflammatory, neuropathic, and musculoskeletal subtypes [40–42], and our primary goal was to test whether being in a state of pain over prolonged periods of time - regardless of diagnosis - was associated with shifts in decision-making. Whether distinct chronic pain conditions show different valuation profiles remains an open question requiring larger, phenotypically characterized samples. A particularly interesting question is whether the valuation shift differs depending on the nature of pain onset - for instance, whether it was triggered by a discrete event such as surgery, injury, or childbirth.

Fifth, task instructions and feedback structure differed slightly from earlier studies using similar paradigms [21], which may have influenced the balance between context-dependent and global value coding. Our sample showed a somewhat attenuated context-dependency effect compared to prior work, which may reflect these design differences or population characteristics.

Finally, our cross-sectional design cannot establish whether the valuation shift precedes or follows pain chronification. The strong association between the weighting parameter and pain duration is consistent with a causal role of chronic exposure in reshaping decision-making, but directionality cannot be confirmed without longitudinal data. Prospective studies following individuals from acute injury through potential chronification are needed to determine whether shifts in individual decision-making profiles predict pain persistence - and whether they could serve as targets for early cognitive intervention.

## Conclusion

Chronic pain is accompanied by a duration-dependent shift in decision-making strategy - away from globally integrated expected value and toward what was learned as locally advantageous in a prior context. This shift provides a computational mechanism for the entrenchment of avoidance behaviors that the fear-avoidance model describes but does not fully specify. The intermediate profile of the acute pain group raises the possibility that this reconfiguration begins early in the pain trajectory, before chronification is established. Longitudinal studies are now needed to determine whether individual decision-making profiles predict who goes on to develop chronic pain - and whether targeting this shift could reduce the behavioral burden of chronification.

## Methods

### Participants

Participants were recruited as part of a large-scale study on chronic pain. A total of 360 individuals enrolled via the online data acquisition platform Prolific. Given reports that approximately 30% of Prolific participants may be inattentive or non-human [43], we implemented strict performance-based exclusion criteria to ensure data quality. Specifically, participants were required to correctly answer several task-related comprehension questions following the instructions before proceeding to the experimental phase. In addition, participants were excluded if they failed to achieve an average accuracy of at least 70% during the final quarter of the learning phase (see Task section below), resulting in the exclusion of 57 participants (16%). A further 64 participants (18%) were excluded due to inconsistencies in their pain reports (see Pain Groups section below). The final sample comprised of 239 participants: 108 in the no pain group (age = 33.74 [SD: 11.34], female/male/undisclosed = 54/54/0), 53 in the acute pain group (age = 36.17 [SD: 13.64], f/m/u = 29/23/1), and 78 in the chronic pain group (age = 40.96 [SD: 12.69], f/m/u = 55/22/1). For behavioral analyses, we excluded trials in which participants responded in under 200 ms or over 5000 ms, resulting in the removal of 1.46% of all trials. Participants received monetary compensation for their participation. The study was approved by the local Ethics Committee (IRB No. 2022003301), and all participants provided informed consent in accordance with the Declaration of Helsinki (2013).

### Demographics & Pain Groups

To assess the current pain status and severity, all participants completed a brief self-report survey modeled after the ICD-11 diagnostic guidance for chronic pain classification. The survey included three items, each rated on a 0–10 numerical scale, assessing (1) pain intensity (“Averaged over the past week, how intense is your pain?”), (2) pain unpleasantness (“Averaged over the past week, how unpleasant is your pain?”), and (3) pain interference (“How much has your pain interfered with your activities over the past week?”). These dimensions correspond to the ICD-11-recommended criteria for coding the severity of chronic pain, which emphasize the multidimensional nature of pain-related impairment, including physical intensity, emotional distress, and disruption of daily functioning [44, 45].

Participants also reported the duration of their current pain experience by selecting one of the following options: “I am not in pain,” “< 2 weeks,” “2–4 weeks,” “1–3 months,” “3–6 months,” “6–12 months,” “1–5 years,” “> 5 years,” or “> 10 years.” Additionally, they indicated whether they had been diagnosed with any pain-related condition from a predefined list (e.g., scoliosis, fibromyalgia; Supplementary Table S10).

Based on their self-reported pain duration, participants were categorized into one of three pain groups: **no pain** (“I am not in pain”), **acute pain** (pain duration ≤ 6 months), and **chronic pain** (pain duration > 6 months). This duration-based definition of chronic pain follows standard clinical and research conventions, including those of the International Association for the Study of Pain (IASP [46]), which defines chronic pain as pain that persists beyond the expected period of tissue healing. This is typically considered three to six months after pain onset. Since some people still heal in the 3-6 months after pain started, we used a conservative cut-off duration of six months for defining the chronic pain group in this study [40, 41, 45].

To ensure internal consistency between pain duration and pain severity, we computed a composite pain score for each participant by averaging their ratings of intensity, unpleasantness, and interference. Participants in the no pain group with a composite score > 2 were excluded, as were participants in the acute or chronic pain groups with a composite score < 2. This procedure resulted in the exclusion of 64 participants from the final sample.

As per the definition, the pain groups differed significantly in their self-reported pain experiences. Robust group effects were observed for pain intensity, unpleasantness, interference, and the composite pain score (*ps* < .0001) (Supplementary Figure S6 and Supplementary Table S9). Post hoc comparisons revealed that participants in the chronic pain group reported significantly higher scores than those in the no pain group across all dimensions (*ps* < .0001; *ds* = 3.45 for intensity, 3.79 for unpleasantness, 3.22 for interference, and 3.80 for the composite score). No significant differences were found between the chronic and acute pain groups (*ps* > .05; *ds*= 0.29, 0.24, 0.08, and 0.22, respectively), indicating the same pain severity levels despite differences in chronicity.

In addition, there was a significant group difference in age (*p* < .0001), with participants in the chronic pain group being older than those in the no pain (*p* < .0001, *d* = 0.61) and acute pain (*p* < .0001, *d* = 0.37) groups. However, no age difference was found between the no pain and acute pain groups (*p* = .4691, *d* = 0.20).

To confirm that observed behavioral effects were not confounded by age or pain severity, we repeated all main behavioral analyses while including each of age and composite pain intensity as covariates. The results remained unchanged, indicating that the group effects observed in learning and decision-making were not attributable to these covariates, but rather to the pain state (acute versus chronic) itself (Supplementary Tables S3 and S11).

### Task

Participants completed a contextual probabilistic two-armed bandit task with a transfer phase, adapted from prior reinforcement learning studies [19–21] (Figure 1a). The task was designed to dissociate two valuation strategies: decision-making based on absolute expected value versus context-specific reinforcement history. The task consisted of two phases: a learning phase and a transfer phase.

#### Learning Phase

During the learning phase, participants made choices between four fixed pairs of abstract visual stimuli (“knights”; i.e., A-B, C-D, E-F, G-H), each distinguishable by a unique chestplate symbol randomly selected from a predefined set of Agathodaimon characters. Each stimulus pair was consistently associated with one of two learning contexts: a reward-seeking context or a punishment-avoidance context. These contexts were signaled in two ways: (1) by the background image on which the pair appeared (either a forest or a desert scene, randomly assigned across participants), and (2) by the identity of the stimulus pair itself. Specifically, the A–B and E–F pairs always appeared in the reward context and involved potential gains, while the C–D and G–H pairs were always presented in the punishment context and involved potential losses. Thus, both the visual background and the stimulus pairing provided distinct cues about the current context.

Within each pair, one stimulus was consistently more advantageous than the other. In the reward context, the high reward rate stimulus (HR, e.g., A) delivered a gain (+10 points) on 75% of trials and zero otherwise, while the low reward rate stimulus (LR, e.g., B) delivered the same gain on only 25% of trials. In the punishment context, the high punishment rate stimulus (HP, e.g., C) resulted in a loss (–10 points) on 75% of trials and zero otherwise, whereas the low punishment rate stimulus (LP, e.g., D) produced the same loss on only 25% of trials. After each choice, participants received feedback on the outcomes of both options, though only the outcome of the chosen stimulus affected their earnings. This learning phase structure required participants to learn which option was relatively better within each pair (Figure 1b,c). Participants were informed that they would receive a monetary performance bonus at the end of the task based on their overall task performance.

Each of the four pairs was presented 24 times in a randomized order, yielding a total of 96 learning trials. On each trial, participants had up to 10 seconds to choose by pressing the ’F’ key (for the left option) or the ’J’ key (for the right). Upon selection, the chosen knight was briefly highlighted, followed by outcome feedback. Feedback was displayed above both options for 2000 ms, with outcome values color-coded to indicate valence: blue for gains, red for losses, and black for neutral outcomes. In addition to numeric feedback, visual cues on the knights (e.g., eye and symbol color) changed to match the outcome valence.

#### Transfer Phase

Following the learning phase, participants entered a transfer phase designed to assess how previously acquired values were generalized to novel decision contexts. In this phase, all eight stimuli from the learning phase (HR, LR, LP, HP from two pairs each) were recombined into every possible pairwise combination. In addition, a novel ninth stimulus—never seen during learning—was introduced and paired once with each of the original eight. This resulted in 36 unique stimulus pairs.

To test for contextual influences of the background, each of these 36 pairs was presented across three background conditions: (1) the background associated with the reward context, (2) the background associated with the punishment context, and (3) a neutral grey background not previously used. This manipulation resulted in 108 unique trials (36 pairs × 3 contexts), each of which was repeated four times in randomized order, yielding a total of 432 transfer trials.

The sequence and timing of each transfer trial mirrored that of the learning phase: participants viewed a pair of stimuli and made a selection using the ’F’ or ’J’ key within a 10-second window. However, no feedback was provided during the transfer phase, ensuring that choices reflected previously learned value representations and choice preferences rather than ongoing learning.

Critically, this design allowed us to disentangle distinct reward-based decision-making strategies, either based on global expected values or context-specific reinforcement learning history. Global expected value-based choices reflect outcome probabilities aggregated across the task, independent of local context. For example, a stimulus with a 25% chance of gain (LR) is objectively more valuable than one with a 25% chance of loss (LP), even though the former was the worse choice, while the latter was the best choice, within their original contexts. Preference for the LR stimulus in this case indicates reliance on global expected value. Context-specific reinforcement history, by contrast, reflects local learning within a fixed stimulus pair and decision context. Stimuli that were consistently the better option within their original pair (i.e., HR and LP) were associated with more frequent positive prediction errors during learning. For example, while LP has a negative absolute expected value, it was the relatively better option in its context (a 75% chance of avoiding loss). It was thus often associated with better-than-expected outcomes. Preference for LP over LR—despite its lower expected value—indicates sensitivity to recent, context-bound reinforcement signals.

By systematically varying pairings and contexts in the absence of feedback, the transfer phase provided a precise behavioral assay of each participant’s decision-making strategy.

#### Instructions and Comprehension Check

Before beginning the main experiment, participants completed a guided instruction phase that familiarized them with the task structure and response procedures (see Supplementary Section Task Instructions). This phase included ten practice trials for each context—reward and punishment—using a separate set of stimuli and background images not used in the main task. These trials demonstrated the core principles of context-dependent learning and probabilistic feedback.

After the practice trials, participants were required to answer four comprehension questions correctly in order to proceed. These questions assessed their understanding of: (1) which keys to use for making responses, (2) how rewards and losses were accrued, (3) how to determine which knight in a pair was better, and (4) how their performance bonus would be calculated. Participants had up to three attempts to answer all questions correctly. If they failed to do so, they were excluded from the study and did not continue to the learning or transfer phases.

#### Task Versions

The data used in this study were collected using two task versions: version A (n = 84) and version B (n = 155). Thus far, we have described version B; however, there were slight differences between version A and version B. In version A, participants received dollar feedback (i.e., +$2, $0, −$2) instead of point feedback, and they selected stimuli using the left and right arrow keys rather than the F and J keys. Furthermore, in version A, participants conducted the experiment as described twice, with a break in between; however, data only from the first implementation were used in our analyses. We confirmed the task version did not affect our findings by repeating all main behavioral analyses with task version as a covariate (Supplementary Tables S3 and S11).

### Statistical Analyses

#### Learning Phase

Performance during the learning phase was quantified using two measures: accuracy and reaction time (RT). Accuracy was defined as the proportion of trials in which the objectively better stimulus was selected, specifically, the high reward (HR) over the low reward (LR) stimulus in the reward context, and the low punishment (LP) over the high punishment (HP) stimulus in the punishment context. Reaction times were recorded as the latency between stimulus onset and response. As RTs exhibited a positive skew, a log transformation was applied prior to analyses.

To examine the temporal dynamics of learning, the 24 presentations of each stimulus pair were divided into four trial bins (hereafter named time): early (trials 1–6), mid-early (7–12), mid-late (13–18), and late (19–24). This binning enabled the time-resolved assessment of changes in accuracy and RT across the learning period.

Statistical analyses were conducted using generalized linear mixed-effects models (GLMMs). For both accuracy and RT, fixed effects included group (no pain, acute pain, chronic pain), context (reward, punishment), and time (trial bins), as well as all two- and three-way interactions. Participant identity was entered as a random effect, allowing the intercept to vary across participants. Accuracy was modeled using a binomial distribution as per equation (1), and RTs were modeled using a gamma distribution with a log link function as per equation (2):

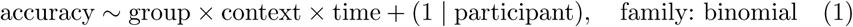

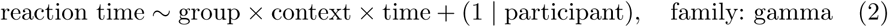

Planned comparisons were conducted to test specific hypotheses regarding group differences. For the main effect of group, two independent-samples t-tests were performed to compare the chronic pain group against the no pain and acute pain groups. To assess the group-by-context interaction, accuracy and RTs were compared between the chronic and no pain groups within each context separately, as well as for the difference between reward and punishment contexts. All additional comparisons were evaluated using Tukey’s Honestly Significant Difference (HSD) post hoc tests.

#### Transfer Phase

Performance during the transfer phase was quantified using two measures: choice rate and RT. Choice rate was defined as the proportion of times a given stimulus was selected when paired with another. Trials in which both stimuli had identical values (e.g., two high reward stimuli) were excluded from analysis, as they provided no informative contrast. RTs were defined as the time, in milliseconds, taken to select one of the two stimuli from their presentation, and again log-transformed.

Choice rates and reaction times were analyzed using GLMMs. Fixed effects included group (no pain, acute pain, chronic pain), stimulus type (HR, LR, LP, HP, Novel [N]), and their interaction. Participant was again modeled as a random intercept. Choice rates were modeled using a Gaussian distribution as per equation (3), and reaction times were modeled using a gamma distribution with a log link function as per equation (4):

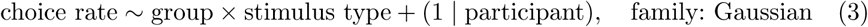

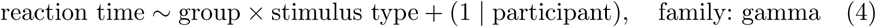

Planned comparisons were conducted to test specific hypotheses regarding group differences in valuation strategy. For both outcome variables, the no pain and acute pain groups were compared to the chronic pain group to detect overall group effects. To investigate the influence of valuation strategy, two stimulus comparisons were of particular interest.

First, the contrast between HR and LP stimuli (HR-LP) was used to assess sensitivity to objective value. Second, the contrast between LR and LP stimuli (LR-LP) allowed us to examine situations in which objective and relative values diverged. Preference for LR would suggest reliance on global value representations, while preference for LP would reflect sensitivity to reinforcement history. Equal preference for both would imply equal integration of both dimensions.

For the group-by-stimulus interaction, planned comparisons were conducted to examine group differences in these two contrasts (HR-LP and LR-LP). Specifically, the no pain, acute pain, and chronic pain groups were compared pairwise to assess whether valuation strategies varied as a function of pain state. These planned comparisons functioned on choice rates specific to these pairs of data—for example, the HR-LP choice rates would be the number of times that the HR stimulus type was chosen when paired with the LP stimulus type minus the number of times that the LP stimulus type was chosen when paired with the HR stimulus type (which would be the complement of the first). All remaining comparisons were evaluated using Tukey’s Honestly Significant Difference (HSD) post hoc tests.

#### Significance

Statistical significance was determined using a threshold of *α < .*05. For all GLMMs, assumptions of linearity, homoskedasticity, and normality of residuals were assessed. Linearity was satisfied in all cases. Although violations of homoskedasticity and normality were observed, these assumptions have been shown to have a limited impact on the validity of GLMMs [47]; therefore, no corrective measures were applied. For independent-samples t-tests, assumptions of normality and homogeneity of variance were evaluated. Where necessary, variables exhibiting non-normal distributions were transformed. When the assumption of equal variances was violated, Welch’s t-tests were used in place of standard t-tests.

### Computational Modeling

Computational modeling was used to quantify the cognitive mechanisms underlying group differences in decision-making. As described earlier, reward-based decision-making can be informed by two factors: an objective dimension based on absolute expected value, and a relative dimension shaped by context-specific reinforcement history. To dissociate the influence of these factors, we fit a set of reinforcement learning models that captured each dimension independently or in combination: a Q-Learning model to represent pure expected value learning, an Actor-Critic model to capture sensitivity to local reinforcement [23], a Relative model [21], a Hybrid model [19, 20], and an Advantage model that all integrate both factors in different ways.

#### Q-Learning Model

To capture decision-making based on absolute expected value, we implemented a Q-Learning model with counterfactual feedback, which updates action values for both the chosen and unchosen option on each trial [22]. In this model, *Q*-values represent the expected value of each action in a given state and are updated according to a delta rule. For the chosen action *α_c_* and unchosen action *α_u_*, the updates are defined as equations (5) and (6):

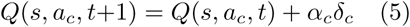

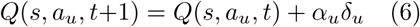

where *α_c_* and *α_u_* are the learning rates for the chosen and unchosen actions, respectively. Prediction errors, *δ_c_* and *δ_u_*, are computed as equations (7) and (8):

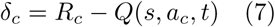

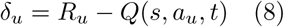

where *R_c_* and *R_u_* are the outcome feedback for the chosen and unchosen action, respectively. To standardize across gain and loss contexts, outcomes were rescaled to −1 (loss), 0 (neutral), and +1 (gain) prior to model fitting.

Action selection was modeled using a softmax choice rule, as per equation (9):

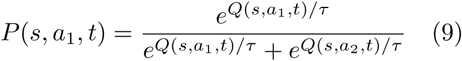

where *τ* is the temperature parameter controlling choice stochasticity. The complete model, therefore, includes three free parameters: *α_c_*, *α_u_*, and *τ* .

#### Actor-Critic

To model decision-making driven by context-specific reinforcement history, we implemented an Actor-Critic model with counterfactual feedback. This architecture reflects the functional organization of basal ganglia circuitry, where action selection is influenced by bottom-up prediction errors relative to the current state value [24].

In this framework, the critic component estimates the value of each state, which is updated based on the outcome of the chosen action, as per equation (10):

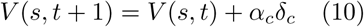

where *α_c_* is the critic learning rate, and *δ_c_* is the prediction error of the chosen action. Prediction errors are computed for the chosen and unchosen actions, which are displayed throughout the learning phase, as equations (11) and (12):

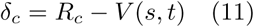

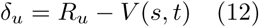

where *R_c_*and *R_u_* are the outcome feedback for the chosen and unchosen actions, respectively. Each action within a state has a *w*-value, which is updated as per equations (13) and (14):

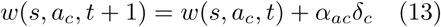

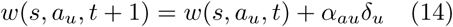

where *α_ac_*is the actor learning rate for the chosen action, and *α_au_* is the actor learning rate for the unchosen action. To prevent unbounded growth of these values, we normalize them as per equation (15):

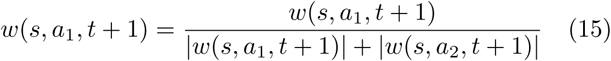

Action selection was modeled using a softmax choice rule as per equation (16):

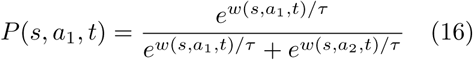

where *τ* is the temperature parameter. This model results in a total of four parameters:

#### Hybrid Model

The Hybrid model captures the combined influence of absolute expected value and relative reinforcement history on decision-making by integrating Q-Learning and Actor-Critic components [19, 20], by computing a weighted mixture of *Q*-value and Actor-Critic weights, indicated in equation (17):

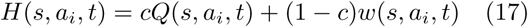

where *Q*(*s, a_i_, t*) reflects the absolute expected value of action *i* in state *s*, learned through the Q-Learning component, and *w*(*s, a_i_, t*) reflects the action preference learned via the Actor-Critic component. The mixing parameter *c* determines the relative contribution of each system: when *c* = 1, the model behaves as a pure Q-Learner; when *c* = 0, it reduces to an Actor-Critic model.

Actions are selected using a softmax decision rule based on *H*-values, as per equation (18):

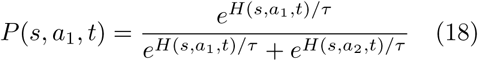

The Hybrid model was initially introduced by Gold et al. [20] and later extended by Geana et al. [19] to include noise and decay parameters. In the present study, we employed a streamlined version of the original Hybrid model, modified to incorporate counterfactual learning rates for both the Q-Learning and Actor-Critic components, and excluding the valence bias parameter. Although we focus here on this specific variant, we systematically evaluated a set of ten Hybrid model variations, including both original formulations and modified versions with or without counterfactual learning, valence bias, and decay terms.

The Hybrid model used in the present study includes seven free parameters: two specific to the Q-Learning component, *α_qc_* and *α_qu_*, three specific to the Actor-Critic component, *α_wc_*, *α_wac_*, and *α_wau_*, and two additional parameters governing decision dynamics—the softmax temperature *τ* and the mixture weight *c*, which determines the relative contribution of each component to the choice behavior.

#### Relative Model

The relative model is a mixture model that learns combined objective-relative values of actions [21]. It is a modified Q-Learning model with the addition of a learned state value, which serves as a reference point when computing prediction errors (see also [48–50]. As with the Q-Learning model, *Q*-values are updated as per equations (5) and (6), which include *α_c_* as the learning rate for the chosen action and *α_u_* as the learning rate for the unchosen action. However, unlike the Q-Learning model, the prediction errors are computed as per equations (19) and (20):

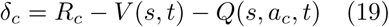

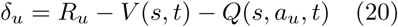

where *R_c_*and *R_u_* are the outcome feedback for the chosen and unchosen actions, respectively. *V* (*s, t*) is the learned state value, which is updated as in equation (21):

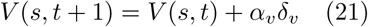

where *α_v_*is the state learning rate. *δ_v_*is the state prediction error, computed as in equation (22):

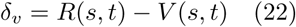

where *R*(*s, t*) is the average reward of each action computed as per equation (23):

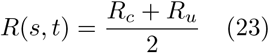

As with the Q-Learning model, action selection uses the softmax function depicted in equation (9), which includes *τ* as the temperature. This model results in a total of four parameters: *α_v_*, *α_c_*, *α_u_*, and *τ* .

#### Advantage Model

Here, we introduce the Advantage model, which is a simplified version of the Relative model [21]. The Advantage model includes the averaged reward of the current state within the prediction error directly, rather than as a learned process. This may be reflected in findings suggesting that striatal dopamine fluctuations encode the integration of a reward prediction error with a counterfactual prediction error, reflecting how much better or worse the current outcome could have been [51]. The prediction errors in the Advantage model are calculated as in equations (24) and (25):

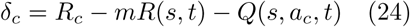

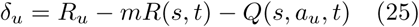

where *m* is a weighting factor determining the influence of reward on the prediction error, and *R*(*s, t*) is the average of the rewards available for the chosen and unchosen option, as described in equation (23). *Q*-value updates remain the same as the Q-Learning and Relative models, as depicted in equations (5) and (6), and so does the action selection function, as per equation (9). This model results in a total of four parameters: *m*, *α_c_*, *α_u_*, and *τ* .

#### Novelty Parameter

During the transfer phase, participants encountered a novel stimulus that had not appeared during the learning phase and was therefore unassociated with any past reinforcement history. In principle, such a stimulus could be treated as neutral, with no expected value. However, behavioral data revealed systematic preferences or aversions toward the novel option, suggesting that participants assigned it a subjective value (see Figure 2). To account for this, all models included an additional free parameter, *η*, which captures the estimated value of the novel stimulus during transfer-based decisions. This parameter allows the model to flexibly accommodate individual tendencies toward novelty seeking (exploration bias) or novelty avoidance.

#### Fitting Procedure

Models were fit to individual participant data by minimizing the negative log-likelihood of observed choices across both the learning and transfer phases. For each participant and model, fitting was repeated ten times with pseudo-randomized initial parameter values to avoid local minima. Initial values were drawn from normal distributions centered on prior estimates derived from the original publications of each model (see Supplementary Table S12), with standard deviations set to one-fifteenth of the parameter’s bounded range. Values falling outside these bounds were clipped to the nearest allowable limit.

Model comparison was conducted using the average Bayesian Information Criterion (BIC) based on each participant *n* and model *m*, calculated as in equation (26):

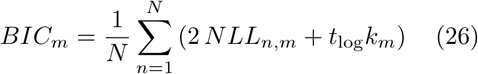

where *NLL_n,m_* is the negative log-likelihood for the corresponding participant *n* and model *m*, *t*_log_ is the log-transformed number of trials for that participant (which is equal across participants), *k_m_* is the number of parameters for the corresponding model, and *N* is the total number of participants.

#### Statistical Analyses of Parameter Estimates

Statistical analyses were performed on the best-fitting model, as determined by BIC. For each model parameter, group differences were assessed using one-way ANOVAs with group (no pain, acute pain, chronic pain) as the between-subject factor. Assumptions of normality and homogeneity of variance were tested; when normality was violated, log-transformations were applied. Homogeneity of variance was generally upheld.

Planned comparisons focused on differences between the no pain group and both the acute and chronic pain groups. For each comparison, independent-samples t-tests were conducted. Welch’s t-tests were used when the assumption of equal variances was violated. Tukey HSD post hoc tests were applied to evaluate all remaining group differences.

We also conducted a series of correlational analyses comparing the different fitted parameters with pain metrics. First, for each of the acute and chronic pain groups, we conducted correlational analyses between the fitted parameters of the winning model (Advantage model: factual learning rate, counterfactual learning rate, temperature, weighting factor, novel value) and the pain metrics (intensity, unpleasantness, interference, composite) using Pearson r correlations. These analyses were not conducted for the no pain group because their range of pain scores was too narrow. Second, we conducted ordinal correlation analyses between the winning model parameter fits and the task duration responses using Spearman Rho correlations. Pain duration responses were coded ordinally as follows: I am not in pain = 0, < 2 weeks = 1, 2–4 weeks = 2, 1–3 months = 3, 3–6 months = 4, 6–12 months = 5, 1–5 years = 6, > 5 years = 7, and > 10 years = 8.

### Data and Code Availability

Data and analysis code for analyses are available within the GitHub repository, avoid learning analysis, while processed data and modeling code for computational modeling are available within the GitHub repository, avoid learning rl models.

## Acknowledgements

This work was supported by the Brainstorm Program at the Robert J. & Nancy D. Carney Institute for Brain Science (FHP) and the COBRE Center for Nervous System Function (NIGMS 5P20GM103645).

## Supplementary Information

### Supplementary Methods

#### Model Validation

To validate the robustness of our computational models and fitting procedures, we conducted a series of standard model checks, following best practices in reinforcement learning research [52]. In addition to the posterior predictive checks, we performed both parameter recovery and model recovery analyses.

For both validations, we generated simulated datasets for the given task structure for each of our 239 participants. Specifically, for each model, we sampled parameter values independently from uniform distributions bounded by each parameter’s plausible range (Supplementary Table S12). Note that high values of temperature would make the models act randomly, making all other parameters obsolete and thus unrecoverable. Due to this, we constrained our model recovery to include temperature bounds of 0.01 to 0.20. These sampled parameters were then used to simulate participant behavior, matching the sequence of contexts, stimulus pairs, and outcomes encountered by each real participant. This procedure resulted in 239 synthetic datasets per model.

These simulated datasets were then submitted to the same model fitting and comparison procedures used in the main analyses, allowing us to assess whether model parameters and model identities could be accurately recovered (Supplementary Figures S7 and S8).

##### Parameter Recovery

To assess whether the models could reliably recover their underlying parameters, we performed parameter recovery analyses on the simulated datasets. For each model, parameters were re-estimated from its corresponding simulated data using the same fitting procedures as in the main manuscript, with two modifications. First, initial parameter values were drawn from uniform distributions constrained to the full range of each parameter, rather than from priors. Second, to improve identifiability, the upper bound for the softmax temperature parameter was reduced from 1.00 to 0.20 during recovery.

Recovered parameters were then compared to the ground-truth values used to generate the simulations, and Pearson correlations were computed as a measure of recovery fidelity. All models demonstrated successful recovery, though to varying degrees (Supplementary Figure S7, Supplementary Table S13). Recovery was highest for the best-fitting model, Advantage, with an average correlation of *r* = .93 across parameters. The Hybrid model showed the weakest recovery overall (*r* = .57), likely due to its greater parameter complexity and non-linear structure. However, even in the Hybrid model, the key parameter indexing the weighting between value systems—the mixing factor—was well recovered (*r* = .89). This suggests that while caution may be warranted when interpreting individual parameters in more complex models, the core mechanisms of interest were robustly estimated.

##### Model Recovery

To verify that each model was selectively identifiable, we conducted model recovery analyses. Specifically, we fit all models to every simulated dataset, using the same fitting procedure as in the parameter recovery analysis. For each simulated dataset, model recovery was assessed using the Bayesian Information Criterion (BIC), with successful recovery defined as the generating model having the lowest BIC among all candidate models. All models showed successful recovery (Supplementary Figure S8).

### Supplemental Results

#### Relative and Hybrid Model Results

As described in the main manuscript, the Advantage model provided the best fit to the empirical data based on model comparison metrics. Accordingly, we focused our main analyses on this model and identified the weighting factor as the key parameter capturing group-level differences in valuation strategy. However, two other mixture models—the Relative and Hybrid models—also showed good overall performance and are widely used in the literature [19–21]. For completeness and comparison, we report the results of these models here in the Supplementary Information. Specifically, we examine model fits, posterior simulations, and group differences in key parameters, and compare these outcomes to those observed in the Advantage model.

##### Relative Model Results

Posterior predictive checks for the Relative model closely replicated the empirical patterns observed in both the learning and transfer phases (Supplementary Figure S9a,b; Supplementary Tables S14, S15). Consistent with the Advantage model, only one parameter—the contextual learning rate—significantly differed across groups (*p* = .0216) (Supplementary Figure S9c). Planned comparisons revealed that this effect was driven by higher contextual learning rates in the chronic pain group compared to the no pain group (*p* = .0062, *d* = 0.41). No significant difference was found between the chronic and acute pain groups (*p* = .1053, *d* = 0.27), and post hoc comparisons indicated no reliable difference between the no pain and acute pain groups (*p* = .2609, *d* = –0.03). These results reinforce the interpretation that chronic pain is associated with a greater reliance on context-dependent relative values.

##### Hybrid Model Results

Posterior predictive checks for the Hybrid model closely reproduced the empirical choice behavior observed across pain groups (Supplementary Figure S10a,b; Supplementary Tables S16, S17). However, unlike the Advantage and Relative models, none of the Hybrid model parameters significantly distinguished between groups (Supplementary Figure S10c). Of particular interest was the mixing parameter, which plays a role analogous to the weighting factor in the Advantage model and the contextual learning rate in the Relative model. This parameter did not differ significantly across groups (*p* = .1766). Planned comparisons revealed no difference between the chronic pain and no pain groups (*p* = .0729, *d* = 0.27), nor between the acute and no pain groups (*p* = .3419, *d* = 0.16).

Although these effects did not reach significance, the downward trend in the mixing parameter from no pain to chronic pain mirrors the group differences captured by the other two models. Given the Hybrid model’s greater complexity and reduced parameter recoverability, these null effects may reflect reduced sensitivity rather than a genuine absence of group differences.

#### Mixture Parameters Capture a Common Latent Mechanism

All three mixture models—the Advantage, Relative, and Hybrid models—include a parameter that modulates the influence of contextual information on value-based learning. In the Advantage model, this is the weighting factor; in the Relative model, it is the contextual learning rate; and in the Hybrid model, it is the mixing parameter. Although these parameters differ in their specific computational implementation, they share a common cognitive function: determining the balance between global expected value and context-specific reinforcement.

The interpretation of higher values differs across models. In the Advantage and Relative models, larger values indicate greater reliance on contextual or locally normalized value representations. In contrast, in the Hybrid model, higher mixing values correspond to increased reliance on global expected value, with reduced contextual influence. To assess whether these parameters capture a shared latent process, we examined their correlations across participants (Supplementary Figure S11). Consistent with their theoretical overlap, we observed strong correlations between all three: Advantage and Relative (*r* = .85, *p* < .0001), Advantage and Hybrid (*r* = –0.59, *p* < .0001), and Relative and Hybrid (*r* = –0.56, *p* < .0001). These findings support the interpretation that all three parameters capture a common valuation mechanism, albeit expressed differently across model architectures.

#### Task Instructions

##### Preamble

In this task, you are picking a team of knights. The knights will look like the ones below. Each knight will have a unique symbol on its chestplate. This symbol will help you identify each knight. You’ll also pick your team of knights from different places, either the desert or the forest. On every turn, you will choose a knight for your team. When you select a knight, it may give you: +$2, +0 dollars, or –$2. Once you’ve selected your knight, their platform and visor will light up to indicate your choice. To help you learn, we will also show you the dollars you could have earned if you had chosen the other knight. Note, you will earn dollars only for the knight you chose. Some knights are better than others. Some will gain dollars while others will avoid losing dollars. Try to earn as many dollars as you can.

##### Learning Phase

During the task, there will be many different knights to choose from. Remember to pay close attention to their symbols on their chestplates. Your job is to try to select the best knight in each pair. Even though you will learn the outcomes for both knights, you will only earn dollars for the knight you choose. Hint: The knights may not always give you dollars, but some knights are better at gaining dollars while other knights are better at avoiding losing dollars. You should try to earn as many dollars as you can, even if it’s not possible to win dollars or avoid losing dollars on every round. At the end of the task, the total number of dollars you’ve earned will be converted into a performance percentage. You can earn up to an additional $4 based on your performance.

##### Transfer Phase

In this next part, you will see the same knights as before, but they will be shown in new pair combinations. Again, your job will be to select the knight you would like to join your team. As you make your choices, you will not receive any feedback after your choice. You should still choose the knight you think is better on each trial. Your choices will still contribute to your performance bonus.

### Supplementary Figures

**Fig. S1.**
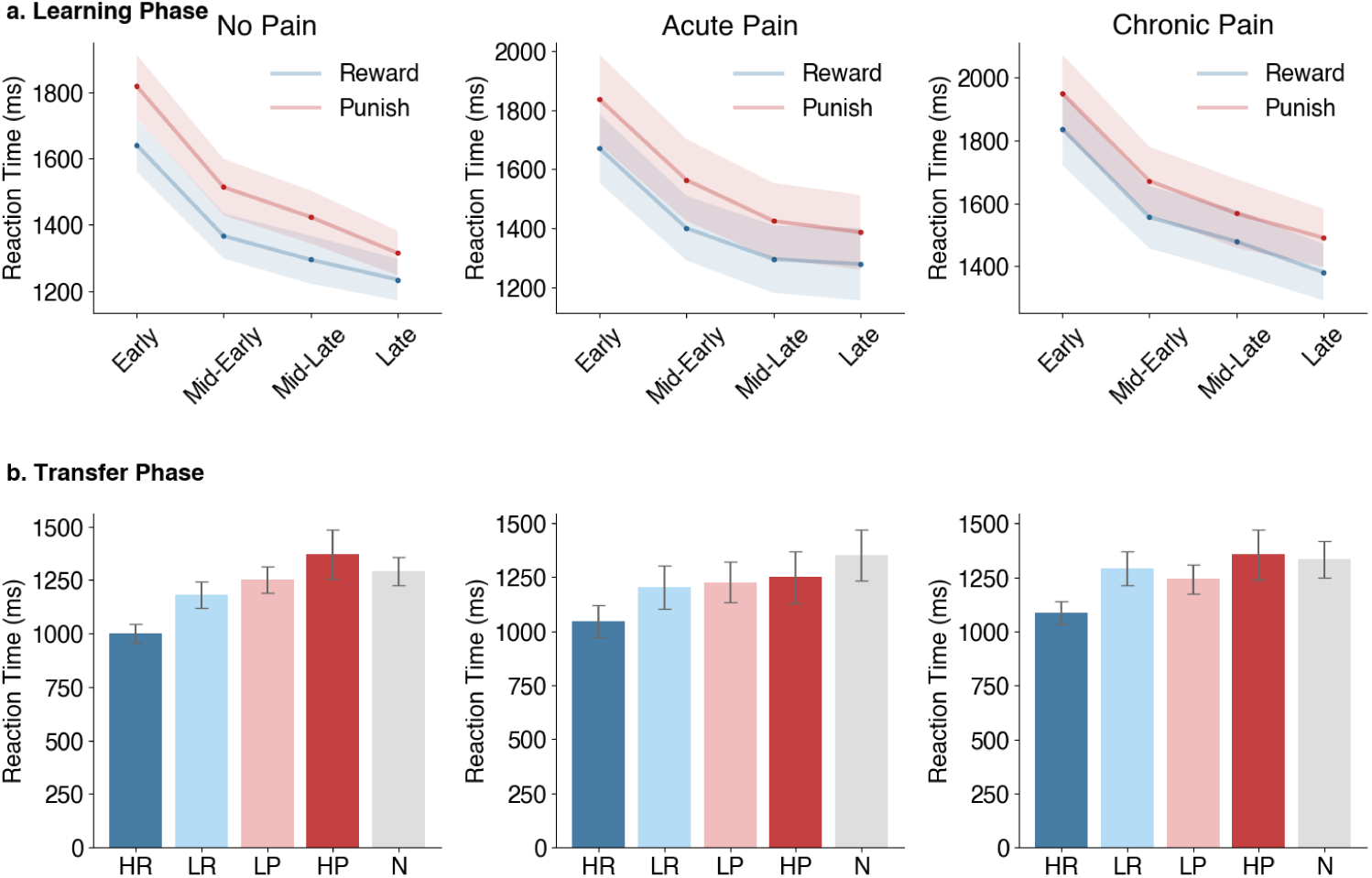
Empirical findings of learning and transfer reaction times. a. Learning Phase: Behavioral reaction times across binned learning trials for the reward and punishment contexts for each group. Shaded regions represent 95% confidence intervals. b. Transfer Phase: Reaction times for each stimulus type during transfer trials for each group. Bar plots show the mean and 95% confidence intervals of the choice rate for each stimulus type across participants within each group. Abbreviations: HR – high reward rate (75% reward), LR – low reward rate (25% reward), LP – low punishment rate (25% loss), HP – high punishment rate (75% loss), N – novel stimulus.

**Fig. S2.**
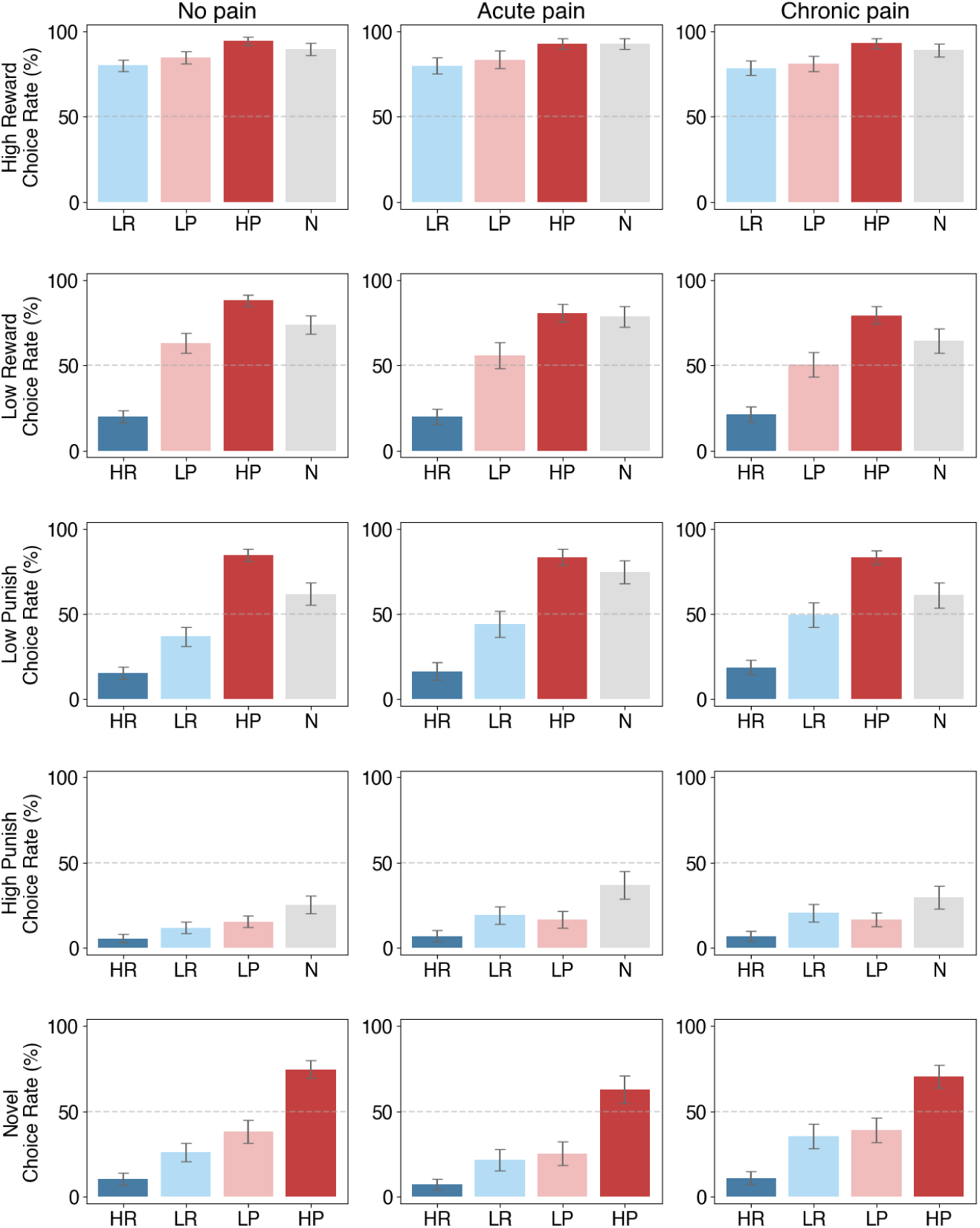
Empirical findings of choice rates. Choice rates for each stimulus type during transfer trials for each group. Data are split where each row represents choice rates anchored to that stimulus type. For example, the first row indicates choice rates of the high reward stimulus type relative to all other stimulus types. This is simply an alternative viewing of the data presented in Figure 2b. Bar plots show the mean and 95% confidence intervals of the choice rate for each stimulus type across participants within each group. Abbreviations: HR – high reward rate (75% reward), LR – low reward rate (25% reward), LP – low punishment rate (25% loss), HP – high punishment rate (75% loss), N – novel stimulus.

**Fig. S3.**
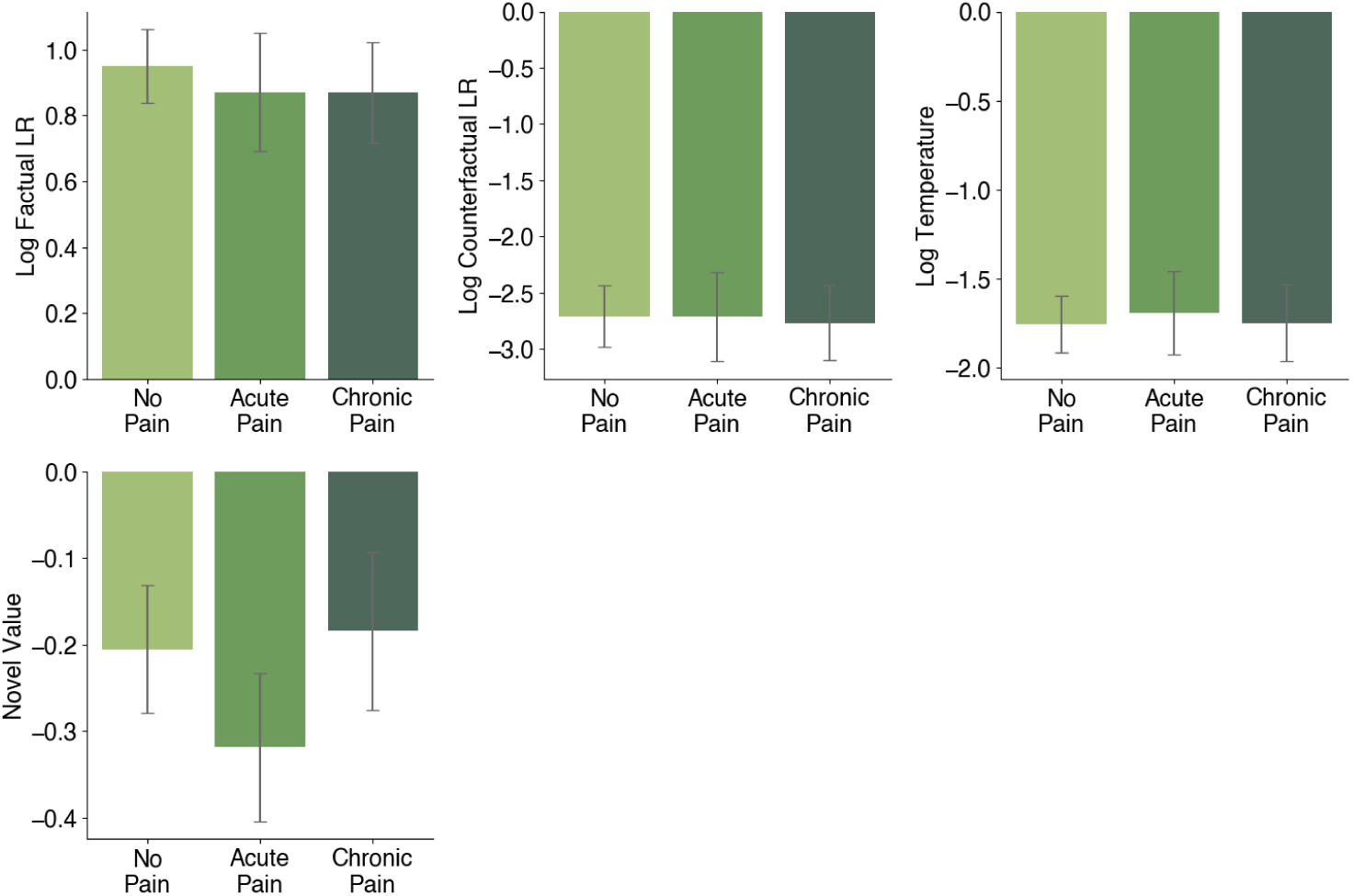
Group effect in the parameters of the Advantage model. Fitted values of the non-significant parameters from the Advantage model. The parameter is displayed as a bar plot, showing the mean and 95% confidence intervals across participants for each group.

**Fig. S4.**
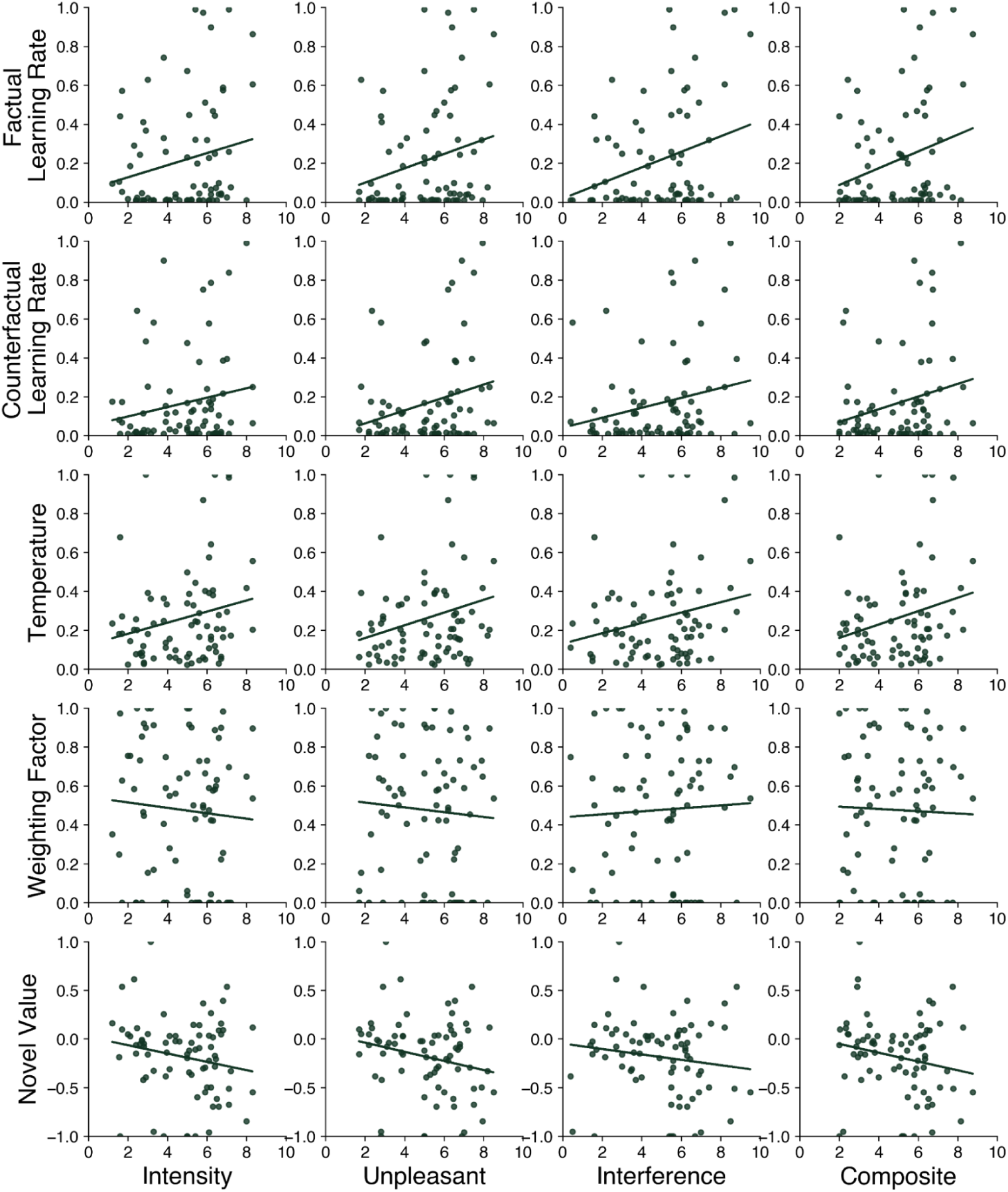
Associations between model parameters and pain severity metrics for chronic pain. Scatterplots with regression lines linking the winning model’s parameter fits with pain severity metrics for the chronic pain group. Associated Pearson *r* correlations can be found in Supplementary Table S8.

**Fig. S5.**
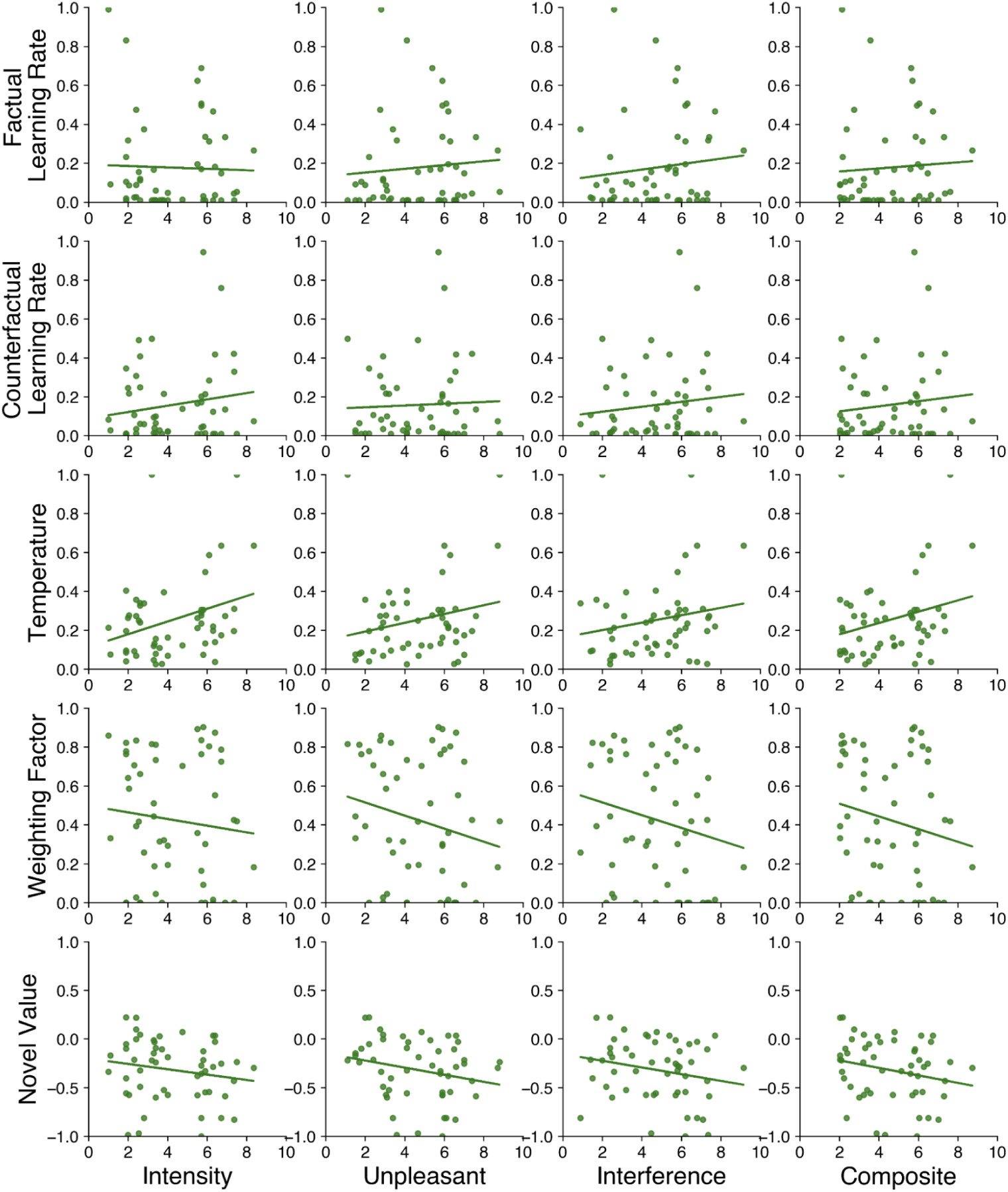
Associations between model parameters and pain severity metrics for acute pain. Scatterplots with regression lines linking the winning model’s parameter fits with pain severity metrics for the acute pain group. Associated Pearson *r* correlations can be found in Supplementary Table S8.

**Fig. S6.**
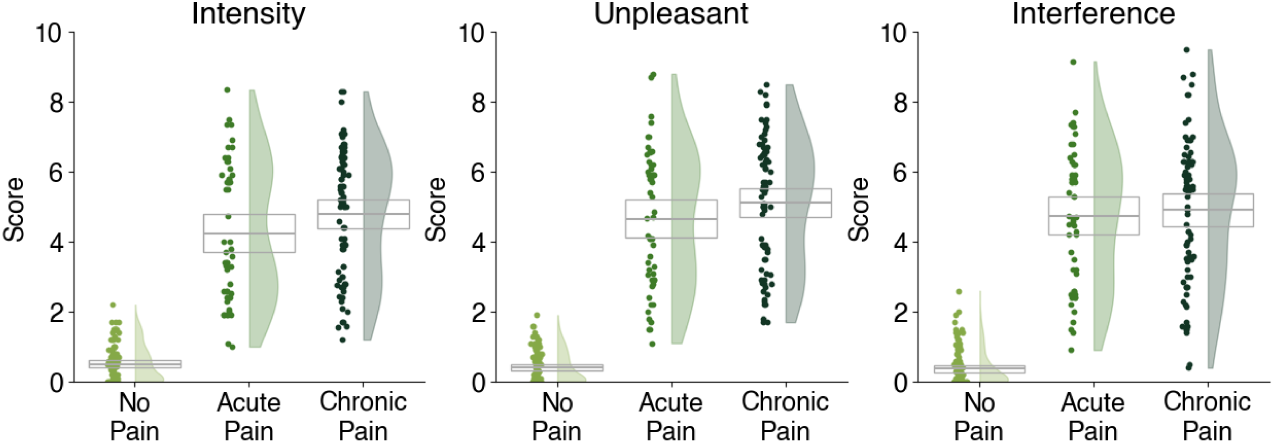
Pain metrics for each group. Boxplots show the mean and 95% confidence intervals of the corresponding metric for each group. Half-violin plots show the distribution of the scores of the corresponding metric for each group. Scatter points show the scores of the corresponding metric for each participant within each group.

**Fig. S7.**
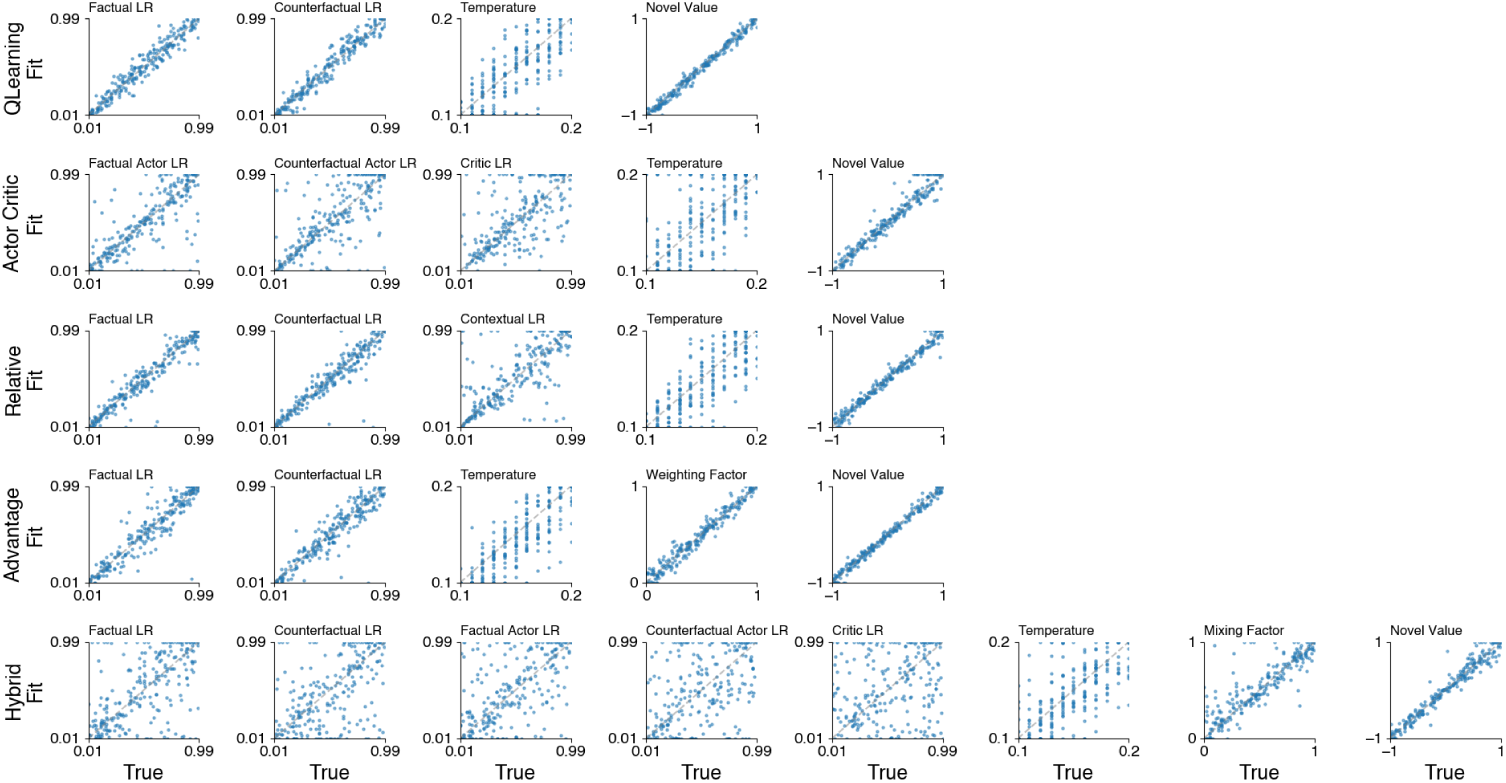
Parameter recovery results. Parameter recovery for each parameter within each model. Data were generated for each model using randomly determined parameters (true values) and then fitted by that model (fit values) to assess the model’s ability to recover parameters. Pearson’s r correlations for each parameter determine the degree to which the parameter was recoverable. These values are presented in Supplementary Table S13. Grey dashed lines indicate a perfect recovery.

**Fig. S8.**
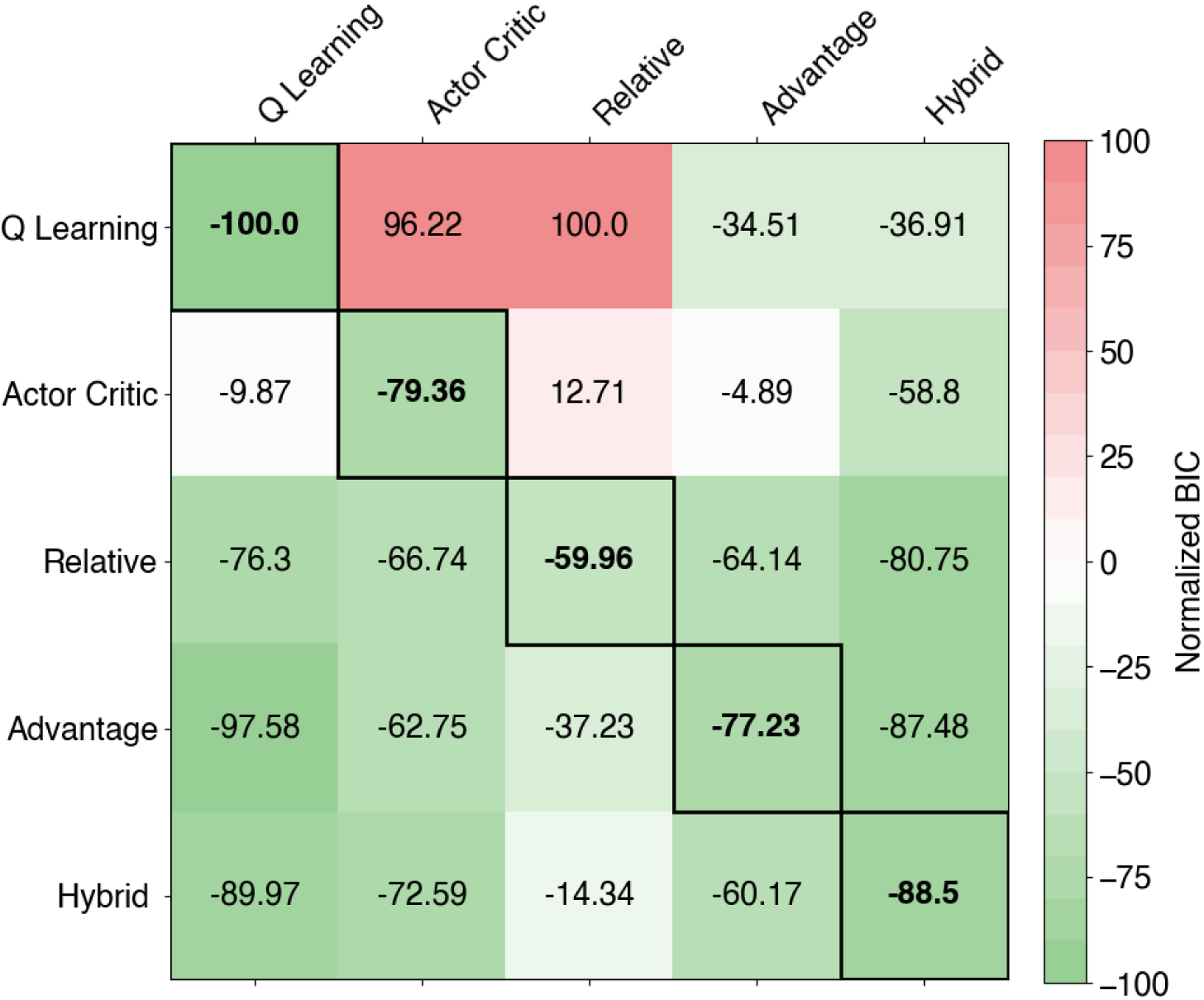
Model recovery results. Model recovery for each model using normalized BIC metrics. Data were generated for each model using randomly determined parameters and then fitted by all models to assess each model’s ability to fit the corresponding model’s data. BIC values are normalized to the range of [0,1], scaled by 200, and then shifted by 100 to center around 0. A BIC of −100 indicates the best fit overall, and a value of +100 indicates the worst fit overall. Within each column, one model fit is highlighted (bolded and bordered), indicating the best-fitted model to explain the generated data. Successful model recovery is when the model best fits itself (as we see here). The values printed are the normalized BIC values.

**Fig. S9.**
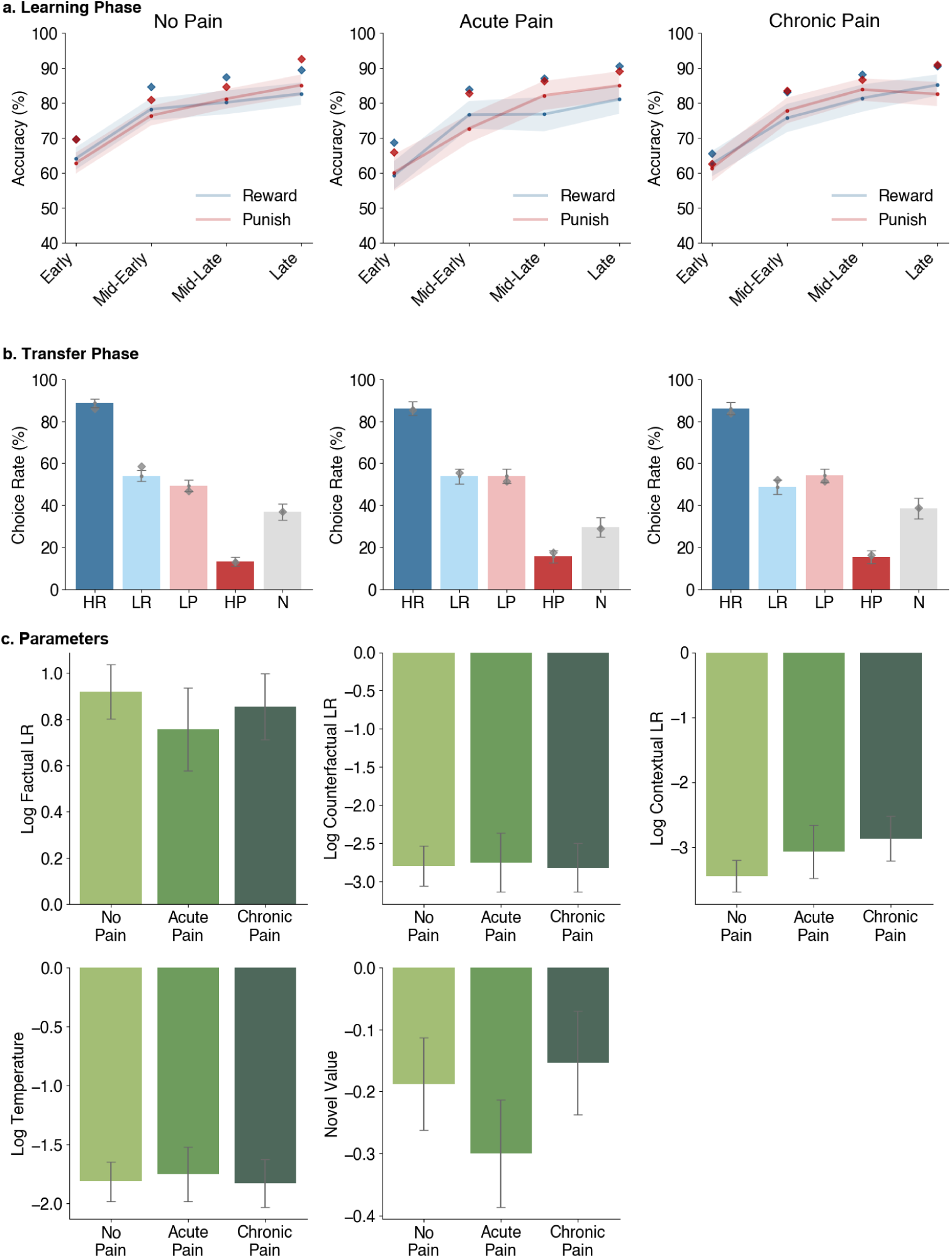
Posterior Predictive checks for the runner-up computational model and fitted parameters. Model simulations of the runner-up model, the Relative model. During the fitting procedure, each participant was fitted ten times, and the best of these runs was selected to determine the participant’s fitted parameters. We then used each model to simulate a single dataset per participant using these fitted parameters. Thus, these data represent the same sample size as our empirical data. a. Learning Phase: Model performance across binned learning trials for the reward and punishment contexts for each group. Shaded regions represent 95% confidence intervals. Blue and red diamonds indicate empirical means of participant accuracy. b. Transfer Phase: Choice rates for each stimulus type during transfer trials for each group. Choice rate is computed as the percentage of times a stimulus type was chosen, given the number of times it was presented. Bar plots show the mean and 95% confidence intervals of the choice rate for each stimulus type across participants within each group. Grey diamonds indicate empirical means of participant choice rates. Abbreviations: HR – high reward rate (75% reward), LR – low reward rate (25% reward), LP – low punishment rate (25% loss), HP – high punishment rate (75% loss), N – novel stimulus. c. Parameters: Fitted parameter values are displayed as a bar plot, showing the mean and 95% confidence intervals across participants for each group.

**Fig. S10.**
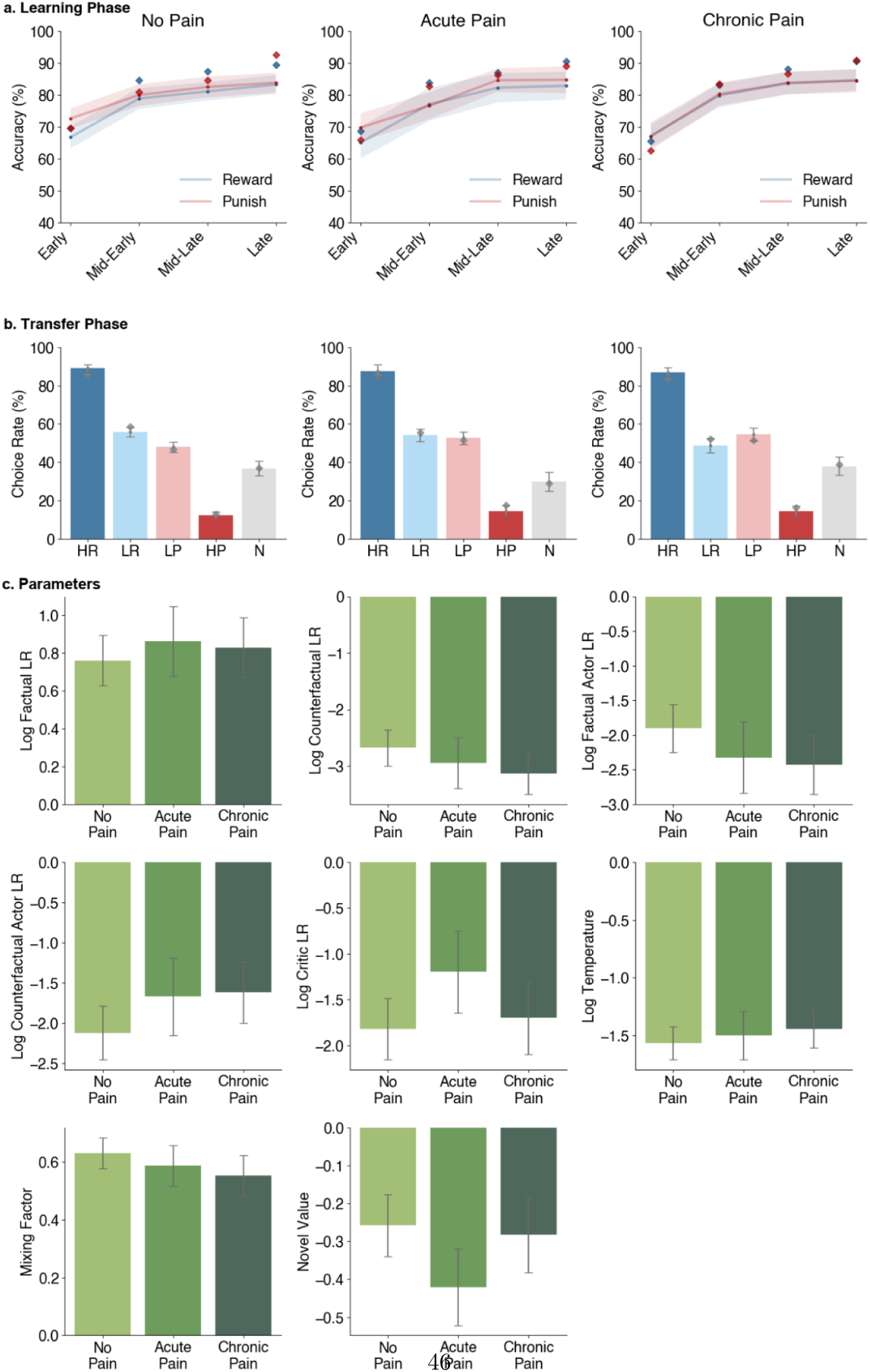
Posterior Predictive checks for the second runner-up computational model and fitted parameters. Model simulations for the second runner-up model, the Hybrid model. During the fitting procedure, each participant was fitted ten times, and the best of these runs was selected to determine the participant’s fitted parameters. We then used each model to simulate a single dataset per participant using these fitted parameters. Thus, these data represent the same sample size as our empirical data. a. Learning Phase: Model performance across binned learning trials for the reward and punishment contexts for each group. Shaded regions represent 95% confidence intervals. Blue and red diamonds indicate empirical means of participant accuracy. b. Transfer Phase: Choice rates for each stimulus type during transfer trials for each group. Choice rate is computed as the percentage of times a stimulus type was chosen, given the number of times it was presented. Bar plots show the mean and 95% confidence intervals of the choice rate for each stimulus type across participants within each group. Grey diamonds indicate empirical means of participant choice rates. Abbreviations: HR – high reward rate (75% reward), LR – low reward rate (25% reward), LP – low punishment rate (25% loss), HP – high punishment rate (75% loss), N – novel stimulus. c. Parameters: Fitted parameter values are displayed as a bar plot, showing the mean and 95% confidence intervals across participants for each group.

**Fig. S11.**
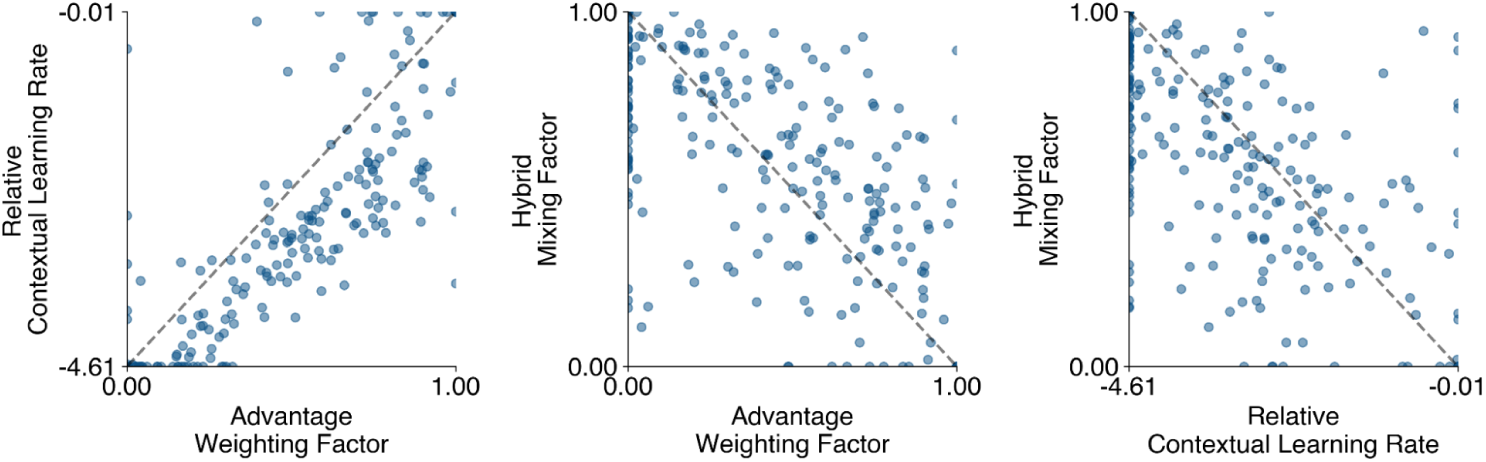
Associations between mixing factors across the mixed computational models. Each of the Advantage, Relative, and Hybrid models includes a parameter that mixes absolute expected values with context-dependent relative values, albeit in different ways. The fit of these values is conceptually related, and these relationships and quantified here. Greater values in the Advantage and Relative models indicate a larger influence of context-dependent relative values, whereas greater values in the Hybrid model indicate a larger influence of absolute expected values. These parameters are highly correlated, suggesting a common underlying mechanism balancing these two valuation strategies.

### Supplementary Tables

**Table S1.** Analysis of accuracy rates during the learning phase. Statistical outcomes of GLMMs (**bold effects**) and planned comparisons (indented and *italicized eflects*) for accuracy rates in the learning phase. Test value represents *χ*^2^ for GLMM effects and t-values for planned comparisons. **Bold p-values** indicate statistically significant effects at *α* = .05.

| Effect | df | Test Value | p-value | Cohen's d |
| --- | --- | --- | --- | --- |
| **Group** | 2 | 0.17 | .9192 |  |
| *Chronic vs No Pain* | 184 | -0.72 | .4730 | -0.11 |
| *Chronic vs Acute* | 129 | -0.26 | .7991 | -0.05 |
| **Context** | 1 | 2.17 | .1406 |  |
| **Time** | 3 | 1117.24 | <.**.0001** |  |
| **Group:Context** | 2 | 0.69 | .7082 |  |
| *Chronic vs No Pain: Reward* | 184 | -0.55 | .5807 | -0.08 |
| *Chronic vs No Pain: Punish* | 184 | -0.57 | .5679 | -0.09 |
| *Chronic vs No Pain: Reward-Punish* | 184 | 0.03 | .9735 | 0.00 |
| **Group:Time** | 6 | 19.32 | **.0037** |  |
| **Context:Time** | 3 | 5.46 | .1409 |  |
| **Group:Context:Time** | 6 | 12.26 | .0565 |  |

**Table S2.** Analysis of reaction times during the learning phase. Statistical outcomes of GLMMs (**bold effects**) and planned comparisons (indented and *italicized eflects*) for reaction times in the learning phase. Test value represents *χ*^2^ for GLMM effects and t-values for planned comparisons. **Bold p-values** indicate statistically significant effects at *α* = .05.

| Effect | df | Test Value | p-value | Cohen's d |
| --- | --- | --- | --- | --- |
| **Group** | 2 | 6.24 | **.0441** |  |
| *Chronic Pain Vs No Pain* | 184 | -3.14 | **.0020** | -0.47 |
| *Chronic Pain Vs Acute Pain* | 129 | -2.31 | **.0224** | -0.41 |
| **Context** | 1 | 270.38 | <.**.0001** |  |
| **Time** | 3 | 1793.61 | <.**.0001** |  |
| **Group:Context** | 2 | 10.37 | **.0056** |  |
| *Chronic Pain Vs No Pain: Reward* | 145 | 3.34 | **.0011** | 0.51 |
| *Chronic Pain Vs No Pain: Punish* | 184 | 2.67 | **.0083** | 0.40 |
| *Chronic Pain Vs No Pain: Reward-Punish* | 184 | 0.81 | .4176 | 0.12 |
| **Group:Time** | 6 | 23.18 | **.0007** |  |
| **Context:Time** | 3 | 3.68 | .2988 |  |
| **Group:Context:Time** | 6 | 3.51 | .7423 |  |

**Table S3.** Analysis of covariates on choice rate during the transfer phase. Age, pain severity, context, and task version were independently added as covariates to the GLMMs assessing choice rates in the transfer phase. These assessments confirm that each covariate does not explain the main findings of our study. Values here represent p-values for each main effect and interaction, as well as the p-value for the corresponding covariate. **Bold p-values** indicate statistically significant effects at *α* = .05.

| Effect | Age | Pain | Context | Task |
| --- | --- | --- | --- | --- |
| Group | .1500 | .2176 | **.0042** | .1504 |
| Stimulus Type | **<.0001** | **<.0001** | **<.0001** | **<.0001** |
| Group:Stimulus Type | **.0002** | **.0002** | **<.0001** | **.0002** |
| Age | .7767 | — | — | — |
| Pain | — | .8222 | — | — |
| Context | — | — | .9873 | — |
| Task | — | — | — | .7279 |

**Table S4.** Analysis of reaction times during the transfer phase. Statistical outcomes of GLMMs (**bold effects**) and planned comparisons (indented and *italicized eflects*) for reaction times in the transfer phase. Test value represents *χ*^2^ for GLMM effects and t-values for planned comparisons. The planned comparisons of the group x stimulus type interaction use slightly different data than the GLMMs and the other planned comparisons. Specifically, these assess reaction times for each pair (e.g., HR vs LP) only when presented in that pair (rather than reaction times across all pairs). **Bold p-values** indicate statistically significant effects at *α* = .05.

| Effect | df | Test Value | p-value | Cohen's d |
| --- | --- | --- | --- | --- |
| **Group** | 2 | 0.23 | .8893 |  |
| *Chronic Pain vs No Pain* | 184 | -0.98 | .3294 | -0.15 |
| *Chronic Pain vs Acute Pain* | 129 | -1.10 | .2720 | -0.20 |
| **Stimulus Type** | 4 | 374.44 | **<.0001** |  |
| *High Reward vs Low Punish* | 238 | -15.08 | **<.0001** | -0.98 |
| *Low Reward vs Low Punish* | 238 | -1.37 | .1732 | -0.09 |
| **Group:Stimulus Type** | 8 | 37.89 | **<.0001** |  |
| *No Pain vs Acute Pain: HR-LP* | 159 | -0.97 | .3361 | -0.18 |
| *No Pain vs Chronic Pain: HR-LP* | 184 | -1.05 | .2966 | -0.17 |
| *Acute Pain vs Chronic Pain: HR-LP* | 129 | 0.11 | .9095 | 0.02 |
| *No Pain vs Acute Pain: LR-LP* | 159 | -1.03 | .3052 | -0.18 |
| *No Pain vs Chronic Pain: LR-LP* | 184 | -1.48 | .1400 | -0.23 |
| *Acute Pain vs Chronic Pain: LR-LP* | 129 | -0.32 | .7528 | -0.06 |

**Table S5.**
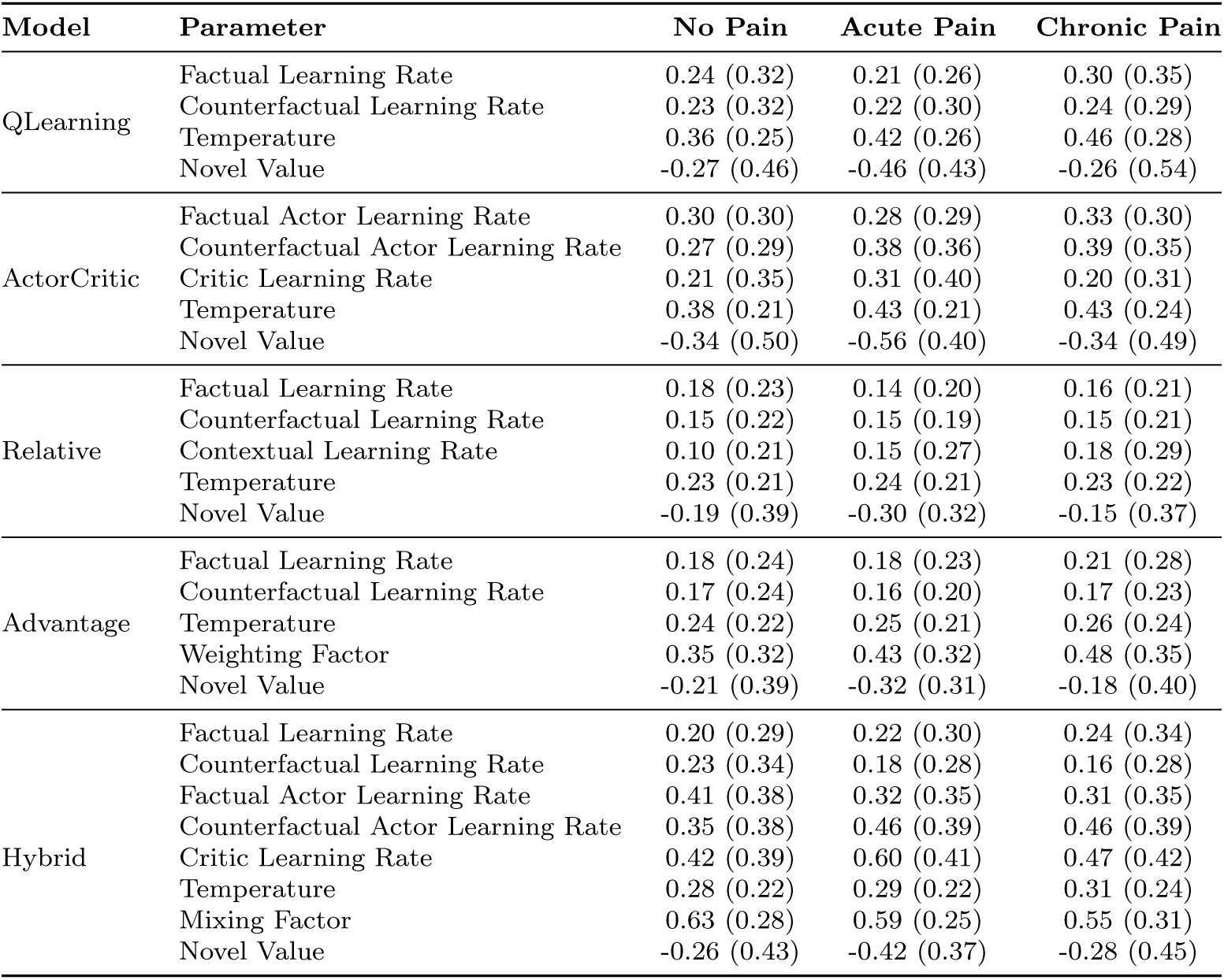
Fitted parameters for each model parameter for each group. Values represent the averaged fitted values, with standard deviations in brackets, for each parameter within each model for each group.

**Table S6.**
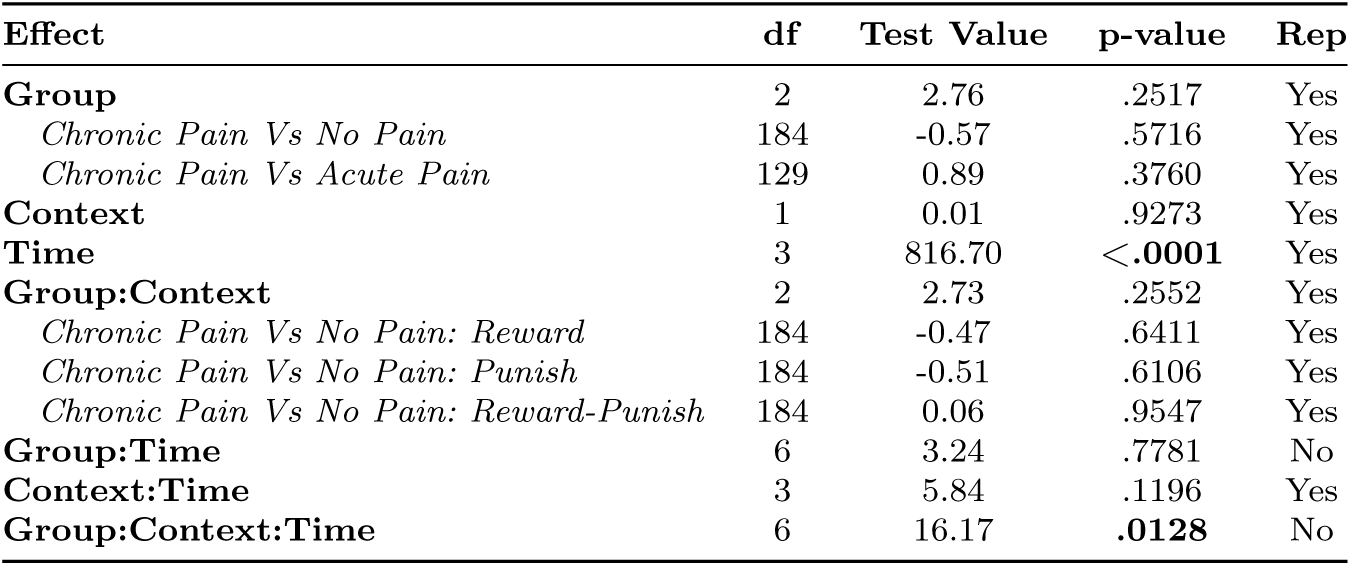
Analysis of model-simulated accuracy rates during the learning phase. Statistical outcomes of GLMMs (**bold effects**) and planned comparisons (indented and *italicized eflects*) for the best-fitting Advantage model simulated accuracy rates in the learning phase. Test value represents *χ*^2^ for GLMM effects and t-values for planned comparisons. **Bold p-values** indicate statistically significant effects at *α* = .05. Rep = replication and indicates whether the model simulations resulted in the same statistical outcome (significant or not significant) as the empirical parallels.

**Table S7.**
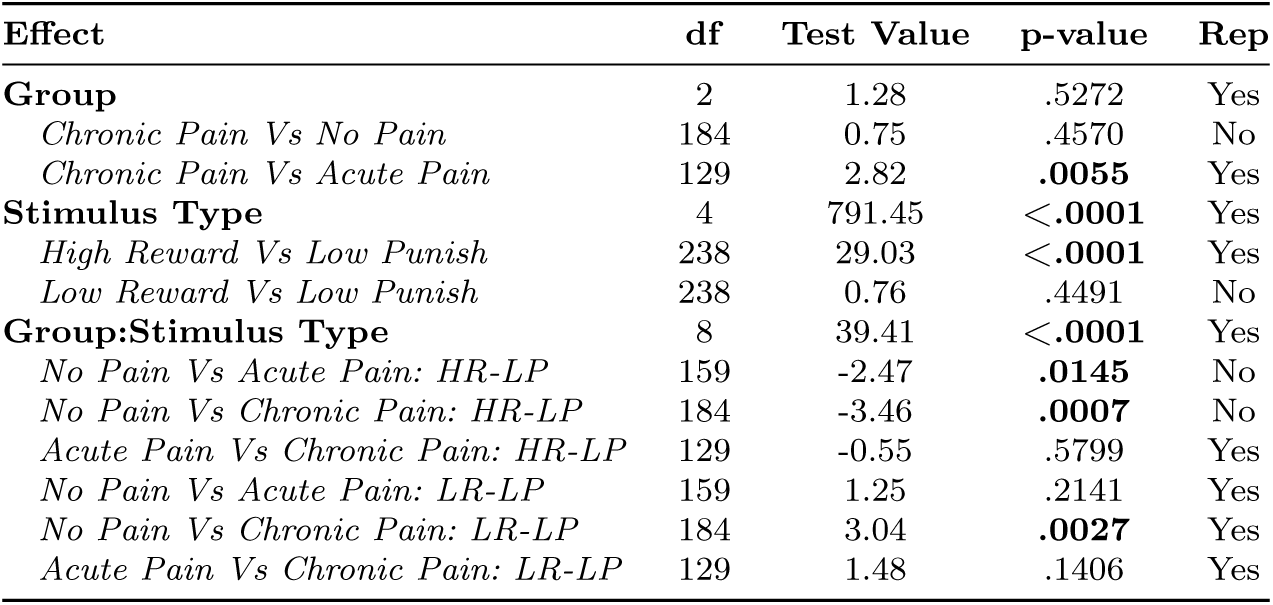
Analysis of model-simulated choice rates during the transfer phase. Statistical outcomes of GLMMs (**bold effects**) and planned comparisons (indented and *italicized eflects*) for the best-fitting Advantage model simulated choice rates in the transfer phase. Test value represents *χ*^2^ for GLMM effects and t-values for planned comparisons. Just as with empirical findings, the planned comparisons of the group x stimulus type interaction use slightly different data than the GLMMs and the other planned comparisons. Specifically, these assess choice rates for each pair (e.g., HR vs LP) only when presented in that pair (rather than choice rates across all pairs). **Bold p-values** indicate statistically significant effects at *α* = .05. Rep = replication and indicates whether the model simulations resulted in the same statistical outcome (significant or not significant) as the empirical parallels.

**Table S8.** Associations between model parameters and pain severity metrics. Results of Pearson *r* correlational analyses linking the winning model’s parameter fits with pain severity metrics. Values indicate Pearson *r* correlations with p-values in brackets. **Bold p-values** indicate statistically significant effects at *α* = .05. LR = Learning Rate.

| Group | Parameter | Intensity | Unpleasant | Interference | Composite |
| --- | --- | --- | --- | --- | --- |
| **No Pain** |  |  |  |  |  |
|  | Factual LR | -.02 (.8558) | -.01 (.9072) | -.12 (.2158) | -.06 (.5396) |
|  | Counterfactual LR | .04 (.6821) | .12 (.2355) | .16 (.1016) | .12 (.2132) |
|  | Temperature | .02 (.8657) | .04 (.6570) | .08 (.4185) | .05 (.5811) |
|  | Weighting Factor | -.10 (.3253) | -.18 (.0674) | -.05 (.6094) | -.12 (.2115) |
|  | Novel Value | .03 (.7238) | .09 (.3403) | .00 (.9840) | .05 (.6259) |
| **Acute Pain** |  |  |  |  |  |
|  | Factual LR | -.03 (.8254) | .08 (.5549) | .12 (.3937) | .06 (.6593) |
|  | Counterfactual LR | .16 (.2485) | .05 (.7446) | .13 (.3694) | .12 (.3920) |
|  | Temperature | **.31 (.0251)** | .21 (.1241) | .18 (.2001) | .25 (.0677) |
|  | Weighting Factor | -.10 (.4600) | -.21 (.1401) | -.20 (.1561) | -.18 (.1893) |
|  | Novel Value | -.17 (.2156) | -.23 (.0950) | -.22 (.1197) | -.22 (.1062) |
| **Chronic Pain** |  |  |  |  |  |
|  | Factual LR | .20 (.0740) | **.24 (.0318)** | **.30 (.0077)** | **.27 (.0156)** |
|  | Counterfactual LR | .19 (.1000) | **.26 (.0212)** | **.23 (.0450)** | **.25 (.0306)** |
|  | Temperature | .21 (.0591) | **.25 (.0282)** | **.23 (.0433)** | **.25 (.0267)** |
|  | Weighting Factor | -.07 (.5265) | -.07 (.5714) | .05 (.6904) | -.03 (.7972) |
|  | Novel Value | -.19 (.0923) | -.21 (.0617) | -.14 (.2123) | -.20 (.0849) |

**Table S9.** Analysis of demographic and pain metrics across groups. Main effects of between-subject ANOVAs (**bold**) and planned comparisons (indented and *italicized*) for each metric assessing group differences. Test value represents F-values for ANOVA effects and t-values for planned comparisons. **Bold p-values** indicate statistically significant effects at *α* = .05.

| Effect | df | Test Value | p-value | Cohen's d |
| --- | --- | --- | --- | --- |
| **Age** | 2, 239 | 7.82 | < .0001 |  |
| *Chronic vs No Pain* | 154 | 4.00 | < .0001 | 0.61 |
| *Chronic vs Acute* | 129 | 2.06 | < .0001 | 0.37 |
| **Pain Intensity** | 2, 239 | 240.50 | < .0001 |  |
| *Chronic vs No Pain* | 87 | 20.25 | < .0001 | 3.45 |
| *Chronic vs Acute* | 129 | 1.65 | .1013 | 0.29 |
| **Pain Unpleasantness** | 2, 239 | 293.46 | < .0001 |  |
| *Chronic vs No Pain* | 84 | 22.11 | < .0001 | 3.79 |
| *Chronic vs Acute* | 129 | 1.34 | .1834 | 0.24 |
| **Pain Interference** | 2, 239 | 246.91 | < .0001 |  |
| *Chronic vs No Pain* | 84 | 18.78 | < .0001 | 3.22 |
| *Chronic vs Acute* | 129 | 0.46 | .6466 | 0.08 |
| **Pain Composite** | 2, 239 | 308.00 | < .0001 |  |
| *Chronic vs No Pain* | 84 | 22.15 | < .0001 | 3.80 |
| *Chronic vs Acute* | 129 | 1.23 | .2215 | 0.22 |

**Table S10.** Counts of body regions where pain originates. Self-reported locations of pain origin in participants from task version B (*n* = 155). These data were not collected for participants from task version A. Participants could indicate multiple regions. The no pain group was excluded as this question is irrelevant without pain. Twenty participants did not respond to this question.

| Region | Acute Pain | Chronic Pain |
| --- | --- | --- |
| Back or neck pain | 5 | 20 |
| Scoliosis | 4 | 6 |
| Osteoarthritis | 1 | 5 |
| Headache | 6 | 4 |
| Migraine | 6 | 4 |
| Irritable bowel syndrome (IBS) | 2 | 3 |
| Temporomandibular joint disorder (TMJ) | 0 | 3 |
| Knee | 1 | 3 |
| Herniation | 0 | 2 |
| Chronic musculoskeletal pain | 0 | 2 |
| Multiple Sclerosis | 0 | 1 |
| Shingles | 0 | 1 |
| Degenerative Disc Disease | 0 | 1 |
| Rheumatoid/osteoarthritis | 0 | 1 |
| Rheumatoid arthritis | 1 | 1 |
| Fibromyalgia | 0 | 1 |
| Post-stroke pain | 0 | 1 |
| Back | 0 | 1 |
| Hip | 0 | 1 |
| Shoulder | 0 | 1 |
| Ankle | 2 | 0 |
| Hand | 1 | 0 |
| Foot | 1 | 0 |
| Other | 5 | 6 |

**Table S11.** Analysis of covariates on accuracy during the learning phase. Age, pain severity, and task version were independently added as covariates to the GLMMs assessing accuracy in the learning phase. These assessments confirm that each covariate does not explain the main findings of our study. Values represent p-values for each main effect and interaction, as well as the p-value for the corresponding covariate. **Bold p-values** indicate statistically significant effects at *α* = .05.

| Effect | Age | Pain | Task |
| --- | --- | --- | --- |
| Group | .9035 | .9171 | .9266 |
| Context | .1407 | .1407 | .1406 |
| Time | **<.0001** | **<.0001** | **<.0001** |
| Group:Context | .7084 | .7084 | .7082 |
| Group:Time | **.0037** | **.0037** | **.0037** |
| Context:Time | .1409 | .1409 | .1409 |
| Group:Context:Time | .0565 | .0565 | .0565 |
| Age | .7635 | — | — |
| Pain | — | .8340 | — |
| Task | — | — | .9524 |

**Table S12.**
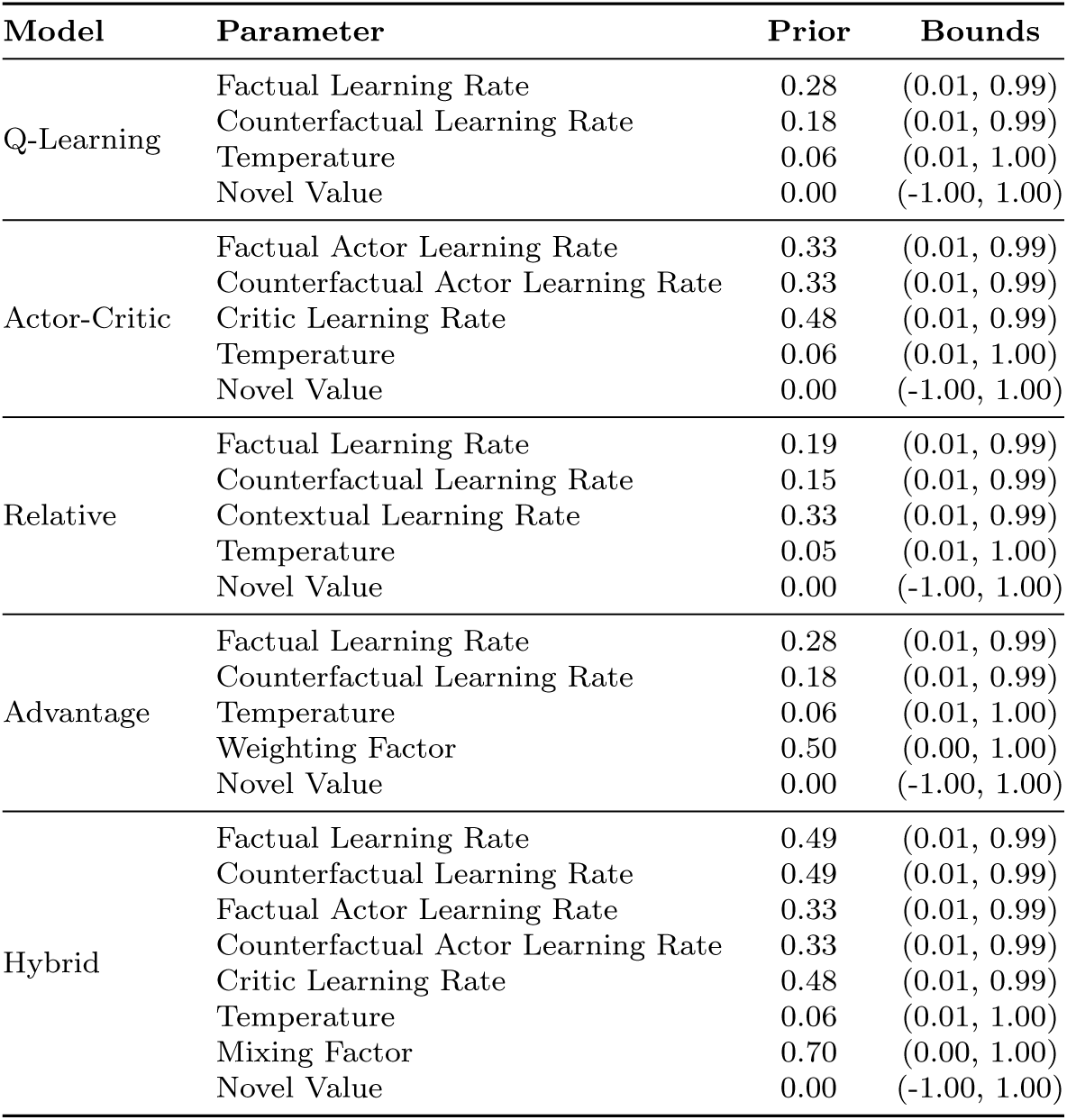
Priors and Bounds for Each Model and Parameter. The priors and bounds used during model fitting of empirical data for each parameter within each model. Priors represent the mean of the normal distribution used as starting points, and bounds constrain each parameter’s valid range.

| Model | Parameter | Prior | Bounds |
| --- | --- | --- | --- |
| Q-Learning | Factual Learning Rate | 0.28 | (0.01, 0.99) |
|  | Counterfactual Learning Rate | 0.18 | (0.01, 0.99) |
|  | Temperature | 0.06 | (0.01, 1.00) |
|  | Novel Value | 0.00 | (-1.00, 1.00) |
| Actor-Critic | Factual Actor Learning Rate | 0.33 | (0.01, 0.99) |
|  | Counterfactual Actor Learning Rate | 0.33 | (0.01, 0.99) |
|  | Critic Learning Rate | 0.48 | (0.01, 0.99) |
|  | Temperature | 0.06 | (0.01, 1.00) |
|  | Novel Value | 0.00 | (-1.00, 1.00) |
| Relative | Factual Learning Rate | 0.19 | (0.01, 0.99) |
|  | Counterfactual Learning Rate | 0.15 | (0.01, 0.99) |
|  | Contextual Learning Rate | 0.33 | (0.01, 0.99) |
|  | Temperature | 0.05 | (0.01, 1.00) |
|  | Novel Value | 0.00 | (-1.00, 1.00) |
| Advantage | Factual Learning Rate | 0.28 | (0.01, 0.99) |
|  | Counterfactual Learning Rate | 0.18 | (0.01, 0.99) |
|  | Temperature | 0.06 | (0.01, 1.00) |
|  | Weighting Factor | 0.50 | (0.00, 1.00) |
|  | Novel Value | 0.00 | (-1.00, 1.00) |
| Hybrid | Factual Learning Rate | 0.49 | (0.01, 0.99) |
|  | Counterfactual Learning Rate | 0.49 | (0.01, 0.99) |
|  | Factual Actor Learning Rate | 0.33 | (0.01, 0.99) |
|  | Counterfactual Actor Learning Rate | 0.33 | (0.01, 0.99) |
|  | Critic Learning Rate | 0.48 | (0.01, 0.99) |
|  | Temperature | 0.06 | (0.01, 1.00) |
|  | Mixing Factor | 0.70 | (0.00, 1.00) |
|  | Novel Value | 0.00 | (-1.00, 1.00) |

**Table S13.** Parameter recovery results. Pearson *r* correlations determining the degree to which each parameter within each model was recoverable. Data were generated for each model using randomly determined parameters and then fitted by that model to assess the model’s ability to recover parameters. Aggregated values indicate the averaged Pearson *r* across parameters within each model. Higher values indicate better recovery.

| Model | Parameter | Pearson $r$ |
| --- | --- | --- |
| Q-Learning | Factual Learning Rate ($\alpha_c$) | 0.97 |
| | Counterfactual Learning Rate ($\alpha_u$) | 0.97 |
| | Temperature ($\tau$) | 0.77 |
| | Novel Value ($\eta$) | 0.99 |
|  | **Aggregated** | **0.93** |
| Actor-Critic | Factual Actor Learning Rate ($\alpha_c$) | 0.80 |
| | Counterfactual Actor Learning Rate ($\alpha_{ac}$) | 0.73 |
| | Critic Learning Rate ($\alpha_{au}$) | 0.63 |
| | Temperature ($\tau$) | 0.62 |
| | Novel Value ($\eta$) | 0.97 |
|  | **Aggregated** | **0.75** |
| Relative | Factual Learning Rate ($\alpha_c$) | 0.95 |
| | Counterfactual Learning Rate ($\alpha_u$) | 0.92 |
| | Contextual Learning Rate ($\alpha_v$) | 0.73 |
| | Temperature ($\tau$) | 0.81 |
| | Novel Value ($\eta$) | 0.99 |
|  | **Aggregated** | **0.88** |
| Advantage | Factual Learning Rate ($\alpha_c$) | 0.93 |
| | Counterfactual Learning Rate ($\alpha_u$) | 0.92 |
| | Temperature ($\tau$) | 0.82 |
| | Weighting Factor ($m$) | 0.98 |
| | Novel Value ($\eta$) | 0.99 |
|  | **Aggregated** | **0.93** |
| Hybrid | Factual Learning Rate ($\alpha_{qc}$) | 0.59 |
| | Counterfactual Learning Rate ($\alpha_{qu}$) | 0.56 |
| | Factual Actor Learning Rate ($\alpha_{wac}$) | 0.56 |
| | Counterfactual Actor Learning Rate ($\alpha_{wau}$) | 0.42 |
| | Critic Learning Rate ($\alpha_{wc}$) | 0.22 |
| | Temperature ($\tau$) | 0.70 |
| | Mixing Factor ($c$) | 0.87 |
| | Novel Value ($\eta$) | 0.98 |
|  | **Aggregated** | **0.61** |

**Table S14.**
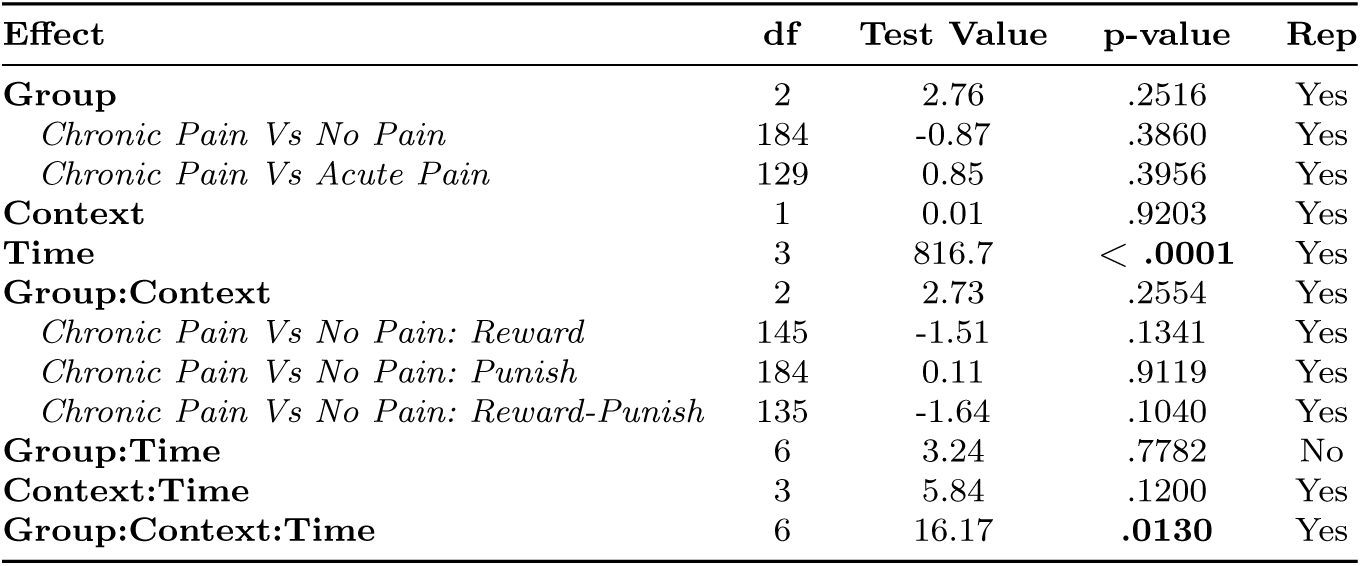
Analysis of model-simulated accuracy rates during the learning phase for the Relative model. Statistical outcomes of GLMMs (**bold effects**) and planned comparisons (indented and *italicized eflects*) for the runner-up Relative model simulated accuracy rates in the learning phase. Test value represents *χ*^2^ for GLMM effects and t-values for planned comparisons. **Bold p-values** indicate statistically significant effects at *α* = .05. *Rep* = replication and indicates whether the model simulations resulted in the same statistical outcome (significant or not significant) as the empirical parallels.

**Table S15.**
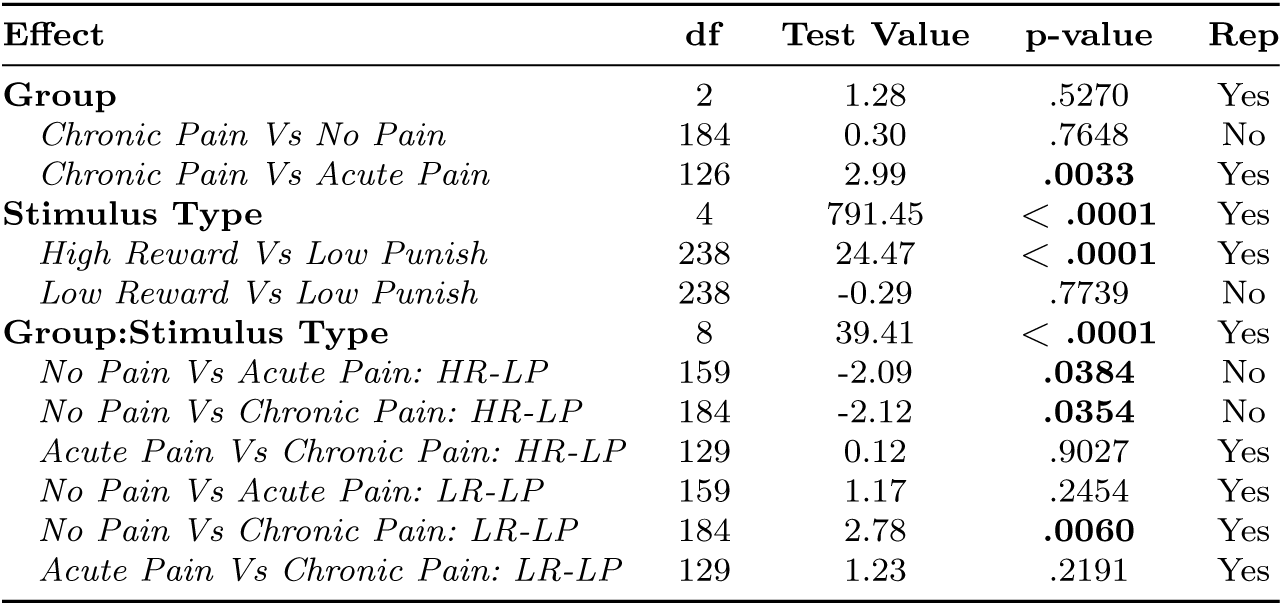
Analysis of model-simulated choice rates during the transfer phase for the Relative model. Statistical outcomes of GLMMs (**bold effects**) and planned comparisons (indented and *italicized eflects*) for the runner-up Relative model simulated choice rates in the transfer phase. Test value represents *χ*^2^ for GLMM effects and t-values for planned comparisons. The planned comparisons of the group x stimulus type interaction use slightly different data than the GLMMs and the other planned comparisons; specifically, these assess choice rates for each pair (e.g., HR vs LP) only when presented in that pair. **Bold p-values** indicate statistically significant effects at *α* = .05. *Rep* indicates whether the model simulations resulted in the same statistical outcome as the empirical parallels.

**Table S16.**
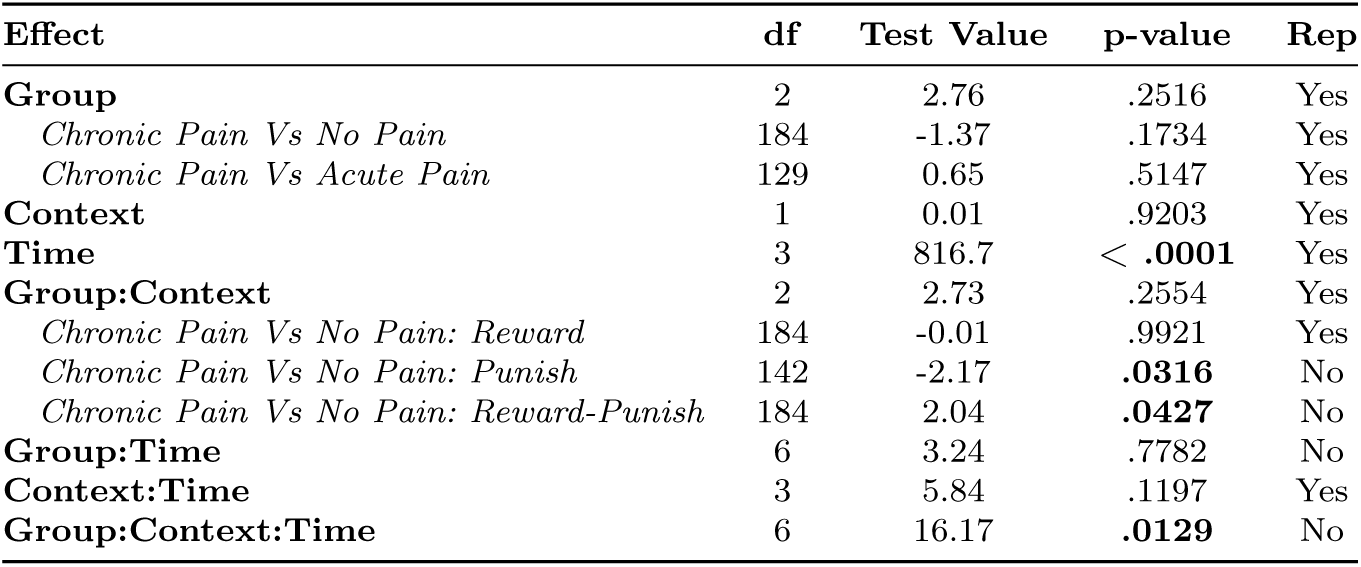
Analysis of model-simulated accuracy rates during the learning phase for the Hybrid model. Statistical outcomes of GLMMs (**bold effects**) and planned comparisons (indented and *italicized effects*) for the second runner-up Hybrid model simulated accuracy rates in the learning phase. Test value represents *χ*^2^ for GLMM effects and t-values for planned comparisons. **Bold p-values** indicate statistically significant effects at *α* = .05. *Rep* indicates whether the model simulations resulted in the same statistical outcome as the empirical parallels.

**Table S17.**
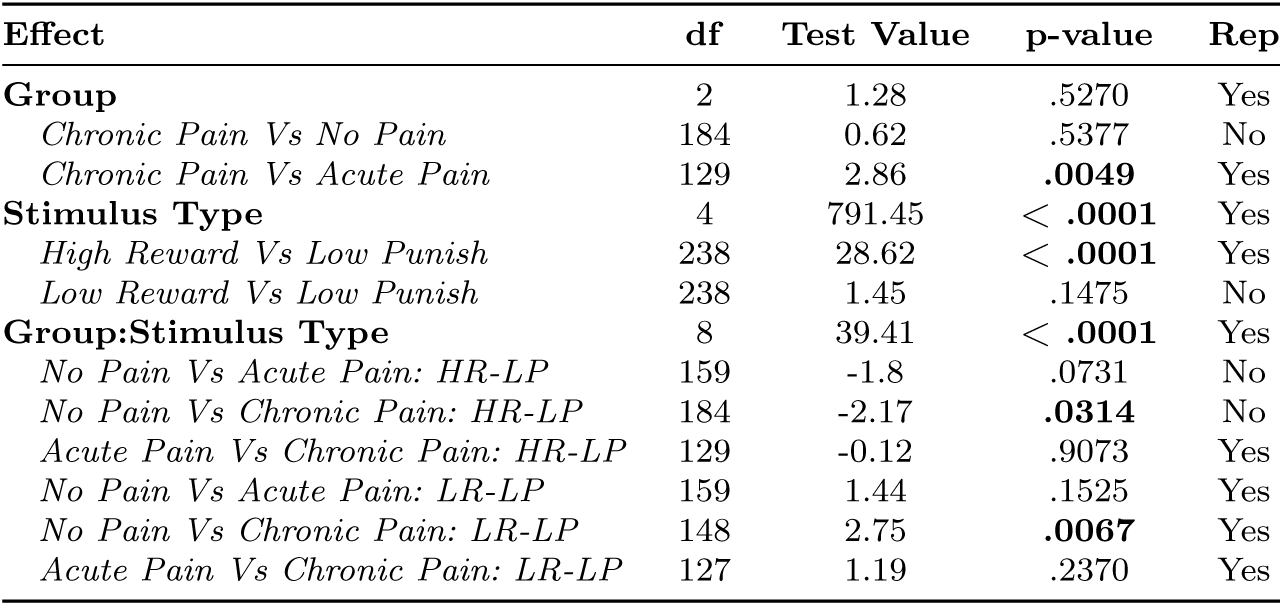
Analysis of model-simulated choice rates during the transfer phase for the Hybrid model. Statistical outcomes of GLMMs (**bold effects**) and planned comparisons (indented and *italicized effects*) for the second runner-up Hybrid model simulated choice rates in the transfer phase. Test value represents *χ*^2^ for GLMM effects and t-values for planned comparisons. Just as with empirical findings, the planned comparisons of the group x stimulus type interaction use slightly different data than the GLMMs and the other planned comparisons; specifically, these assess choice rates for each pair (e.g., HR vs LP) only when presented in that pair (rather than choice rates across all pairs). **Bold p-values** indicate statistically significant effects at *α* = .05. *Rep* = replication and indicates whether the model simulations resulted in the same statistical outcome (significant or not significant) as the empirical parallels.

